# Phosphorylation motif dictates GPCR C-terminal domain conformation and arrestin interaction

**DOI:** 10.1101/2023.02.23.529712

**Authors:** Myriam Guillien, Assia Mouhand, Amin Sagar, Aurélie Fournet, Frédéric Allemand, Glaécia A. N. Pereira, Aurélien Thureau, Pau Bernadó, Jean-Louis Banères, Nathalie Sibille

## Abstract

Arrestin dependent G protein-coupled receptor (GPCR) signaling pathway is regulated by the phosphorylation state of GPCR’s C-terminal domain, but the molecular bases of arrestin:receptor interaction are to be further illuminated. Here we investigated the impact of phosphorylation on the conformational features of the C-terminal region from three Rhodopsin-like GPCRs, the vasopressin V2 Receptor (V2R), the Growth Hormone Secretagogue or ghrelin Receptor type 1a (GHSR) and the β2-Adernergic Receptor (β2AR). Using phosphomimetic variants, we identified pre-formed secondary structure elements, or short linear motif (SLiMs), that undergo specific conformational transitions upon phosphorylation. Of importance, such conformational transition favors arrestin-2 binding. Hence, our results suggest a model in which the cellular signaling specificity of GPCRs is encoded in the phosphorylation-dependent structuration of the C-terminal regions, which will subsequently modulate arrestin conformation and therefore GPCR:arrestin signaling outcomes.

## Introduction

Arrestins are key regulators of G Protein-Coupled Receptor (GPCR) signaling and are involved in three main functions: desensitization, internalization and signaling linked to other scaffolding proteins (Ahn et al., 2020). How these distinct functions are specifically triggered upon GPCR activation remains an open question. A decade ago, it was proposed that the outcome of arrestin signaling depends on the phosphorylation state of active receptors, leading to the so-called phospho-barcode model (Nobles et al., 2011a). This model states that, upon activation by their ligand, GPCRs are phosphorylated at different sites on their C-terminal domain (GPCR-Cter), impacting their interaction with arrestin, in turn affecting arrestin conformation and, ultimately, the intracellular signal triggered by arrestin. Later, atomic-resolution structures of GPCR:arrestin complexes revealed different organization of arrestins, supporting the phospho-barcode model (He et al., 2021a; Huang et al., 2020; Staus et al., 2020; X. Edward Zhou et al., 2017). Additionally, based on the crystal structure of the rhodopsin:arrestin complex, Zhou and co-workers proposed a set of phosphorylation motifs required for the high-affinity binding to arrestin (X. Edward Zhou et al., 2017). These phosphorylation motifs correspond to a long p[S/T]XXp[S/T]XXp[S/T]/E/D motif or a short p[S/T]Xp[S/T]XXp[S/T]/E/D motif, where p[S/T] represents a phospho-serine or -threonine, X is any amino acid except proline in the second XX occurrence. These motifs seem to be conserved across the major GPCR subfamilies despite the very low sequence conservation of GPCR C-terminal domains. Among GPCR:arrestin complexes, a few of them revealed that negatively charged residues and phosphates of GPCR-Cter make a network of electrostatic interactions with conserved basic residues of arrestin pockets (He et al., 2021b; Min et al., 2020; X Edward Zhou et al., 2017). Consequently, it was proposed that these complementary interactions induce conformational changes in arrestin and determine its specific function (He et al., 2021b). This study also supported the existence of a functional phosphorylation barcode system. However, there is still a lack of experimental data explaining how GRK phosphorylation affects the C-terminal domain of GPCRs and leads to different interactions with arrestin.

Interestingly, the GPCR C-terminal domain presents all the structural, dynamic and functional characteristics of an intrinsically disordered region (IDR) (Dunker et al., 2001; Guillien et al., 2020; Uversky et al., 2000; Zhou et al., 2018). Intrinsically disordered proteins (IDPs) or IDRs are highly flexible with a low content of secondary structure, whose specific role in fundamental signaling and regulation processes started to be revealed two decades ago (Kriwacki et al., 1996; Olsen et al., 2017; Wright and Dyson, 2015). We have previously described the transient secondary structure contents of three truncated GPCR C-terminal domains by Nuclear Magnetic Resonance (NMR): the vasopressin V2 Receptor (V2R) (from residue 343 to 371), the Growth Hormone Secretagogue or ghrelin Receptor type 1a (GHSR) (from residue 339 to 366) and the β2-Adernergic Receptor (β2AR) (from residue 342 to 413) (Guillien et al., 2022). Only structures of class B receptors, which interact more strongly with arrestin, or chimeric class A receptors with the C-terminus of V2R (V2Rpp), have been reported in the literature until now (Bous et al., 2022; Huang et al., 2020; Lee et al., 2020; Staus et al., 2020; Yin et al., 2019). In our study, we provide structural information of two class A receptors (GHSR and β2AR), which interact transiently with arrestin. To this aim, we compare these basal non-phosphorylated conformations (wt) to those of the variants mimicking GRK phosphorylation pattern (pm). Sites selectively phosphorylated by GRKs have been described for the three receptors: V2R (Bous et al., 2022; Nobles et al., 2007), GHSR (Bouzo-Lorenzo et al., 2016a), and β2AR (Nobles et al., 2011b). Hence, we designed phosphomimetic variants of these truncated C-terminal regions by replacing all the potential GRK2/3 and GRK5/6 phosphorylation sites (Bouzo-Lorenzo et al., 2016b; Nobles et al., 2011b, 2007) with glutamic acids (Fig. 1a). The use of phosphomimics have been widely used in protein science to decipher the structural and functional role of phosphorylation and avoid chemically heterogenous samples (Bibow et al., 2011; Fischer et al., 2009; Pearlman et al., 2011). Using this strategy, we were able to observe important conformational changes between the non-phosphorylated (wt) and the phosphomimetic (pm) forms. Interestingly, the β-strand conformation appearing in the phosphomimetic variant was found to form specific interactions with arrestin. Moreover, these highlighted binding regions contained motifs described as putative phosphorylation motif, in line with the above-mentioned models of interaction. Altogether, our results suggest that GRK phosphorylation induces the formation of a β-strand in the C-terminal domains of GPCRs that may serve as a general mechanism by which arrestins recognize the phosphorylated carboxy-terminal domains of receptors.

**Fig. 1.**
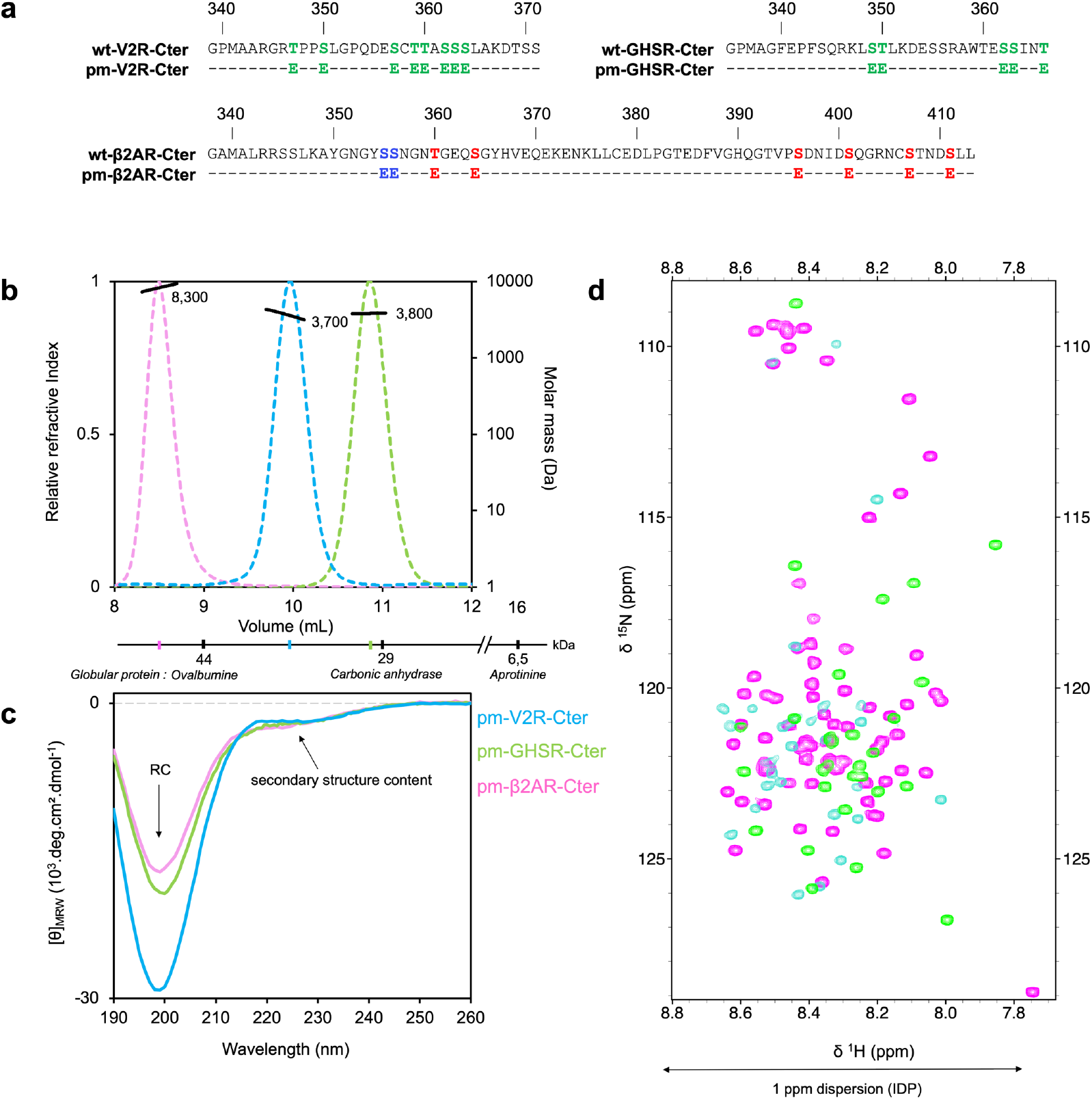
V2R-Cter (blue), GHSR-Cter (green) and β2AR-Cter (pink) phosphomimetic (pm) variants are monomeric and disordered with transient secondary structures. The color code will be maintained for all panels and Figures. **a** Sequence of GPCR-Cters in their non-phosphorylated (wt) and phosphomimetic (pm) form. Phosphorylated sites by GRK2 (red), GRK6 (blue) or any GRKs (green) from the literature for β2AR (Nobles et al., 2011b), GHSR (Bouzo-Lorenzo et al., 2016b), and V2R (Nobles et al., 2007) are indicated in the wt sequence. These sites were mutated to glutamic acid (E) in the pm variants. **b** SEC-MALS profiles (colored dashed lines) of pm variants. Elution volume of GPCR-Cters are smaller than that of globular proteins of the same weight, which is characteristic of elongated/disordered proteins. As a comparison, elution volumes of globular Ovalbumine, Carbonic anhydrase and Aprotinine proteins are indicated under the volume axis. **c** Far-UV Circular Dichroism (CD) spectra (colored lines) of pm variants. **d** ^15^N-HSQC spectra overlaid of pm variants.

## Results

### Structural characterization of the phosphomimetic forms of the three GPCR C-terminal domains

#### Conformational changes between the non-phosphorylated (wt-GPCR-Cter) and phosphomimetic (pm-GPCR-Cter) forms

In order to study the effect of GRK phosphorylation on the conformation of the C-terminal domains of GPCRs (GPCR-Cters), we used three different receptors, i.e., the vasopressin V2 Receptor (V2R), the ghrelin receptor (GHSR) and the β2-Adernergic Receptor (β2AR), as models. These receptors belong to the A (β2AR, GHSR) and B (V2R) classes of GPCRs with regard to interaction with arrestins, where class A interact transiently and class B more tightly (Oakley et al., 2000a). These GPCR-Cter were produced in *E. coli* as phosphomimetic variants (pm-GPCR-Cter). The phosphomimetic (pm) variants were designed by replacing all the serines and threonines phosphorylated by GRK2/3 and 5/6 reported in the literature (Bouzo-Lorenzo et al., 2016b; Nobles et al., 2011b, 2007) with glutamic acids (Fig. 1a). A combined Size Exclusion Chromatography-Multi Angle Light Scattering (SEC-MALS), Circular Dichroism (CD) and Nuclear Magnetic Resonance (NMR) analysis showed that these pm variants are monomeric and disordered domains with transient secondary structures (Fig. 1), as is the case of their non-phosphorylated (wt-GPCR-Cter) forms (Guillien et al., 2022). Indeed, SEC-MALS profiles of pm variants (Fig. 1b) revealed a single peak for each GPCR-Cter at a volume that corresponds to the volume of standard proteins with a molecular mass larger than 13 kDa. This behavior on a SEC column is typical of disordered protein (Uversky, 2012). In contrast, the masses derived from MALS analysis (solid black line) are 3.7 kDa (± 2.4 %) for V2R-Cter, 3.8 kDa (± 1.8 %) for GHSR-Cter and 8.3 kDa (± 0.8 %) for β2AR-Cter (Fig. 1b), which is in agreement with their monomeric molecular weight of 3.6, 3.7 and 8.5 kDa, respectively. This was confirmed by mass spectrometry (MS).

The far-UV Circular Dichroism (CD) spectra of the pm variants present a minimum around 200 nm and a shoulder at 220 nm characteristic of disorder proteins with transient secondary structure contents (Woody, 1996) (Fig. 1c). Moreover, ^15^N-HSQC spectra of pm-V2R-Cter, pm-GHSR-Cter and pm-β2AR-Cter showed a low proton spectral dispersion, typical of IDPs (Sibille and Bernadó, 2012) (Fig. 1d). These observations were further confirmed by Small Angle X-ray Scattering (SAXS) where their Kratky plots displayed the typical profile of an IDP with no clear maximum and a monotonous increase along the momentum transfer range (Fig. S1) (Tiago N. Cordeiro et al., 2017; Tiago N Cordeiro et al., 2017; Sibille and Bernadó, 2012).

To localize and identify secondary structures (Senicourt et al., 2021), we then performed an NMR study. The backbone assignments were first performed for the three phosphomimetic variants (pm) (Fig. S2a-b) (BMRB accession codes: **<u>51319</u>, <u>51328</u>** and **<u>51330</u>** for pm-β2AR-Cter, pm-GHSR-Cter and pm-V2R-Cter respectively). We then used NMR to assess the structural features of the C-terminal domains in their phosphomimetic (pm) form and compared them to their non-phosphorylated (wt) form (Guillien et al., 2022). C-terminal domains were not affected by the protein concentration, as assessed by NMR (Table S1, Fig. S2c).

#### pm-V2R-Cter

the V2R-Cter phosphomimetic variant presented a large stretch with negative ^13^C Secondary Chemical Shift (SCS) values (Fig. 2b), from residues 356 to 364 (called V2-1), suggesting an β-strand conformation in this region. Interestingly, we showed previously that V2-1 contained a helical conformation in the non-phosphorylated form (wt-V2R-Cter) (Guillien et al., 2022). This loss of helical content between the wt and the pm variant was also substantiated by the decrease of percentage of ^3^JHNHA scalar coupling bellow 6 Hz (Fig. S3a), with the majority of scalar couplings remaining in the disordered range between 6 and 8 Hz. Another difference in the SCS profile appeared for residues 346-347 where the pm variant presented highly positive values compared to the wt peptide, certainly related to the presence of a turn. In terms of dynamics, V2-1 relaxation rates R1 and R2 displayed lower values for the variant compared to the wild type (Fig. S4c-d). This could be explained by the fact that the β-strand conformation in pm-V2R-Cter is structurally less constrained than the α-helix in wt-V2R-Cter, even though this change in dynamics was not observed in heteronuclear ^15^N{^1^H}-NOE (Figs. 2c and S4b). Residual Dipolar Couplings (RDC) also indicated the presence of a β-strand conformation in V2-1 region of the variant. The RDCs profile exhibited more negative values in the V2-1 region of the phosphomimetic (pm) variant compared to the non-phosphorylated (wt) form (Figs. 2d and S5), confirming the β-strand conformation in this V2-1 region.

**Fig. 2.**
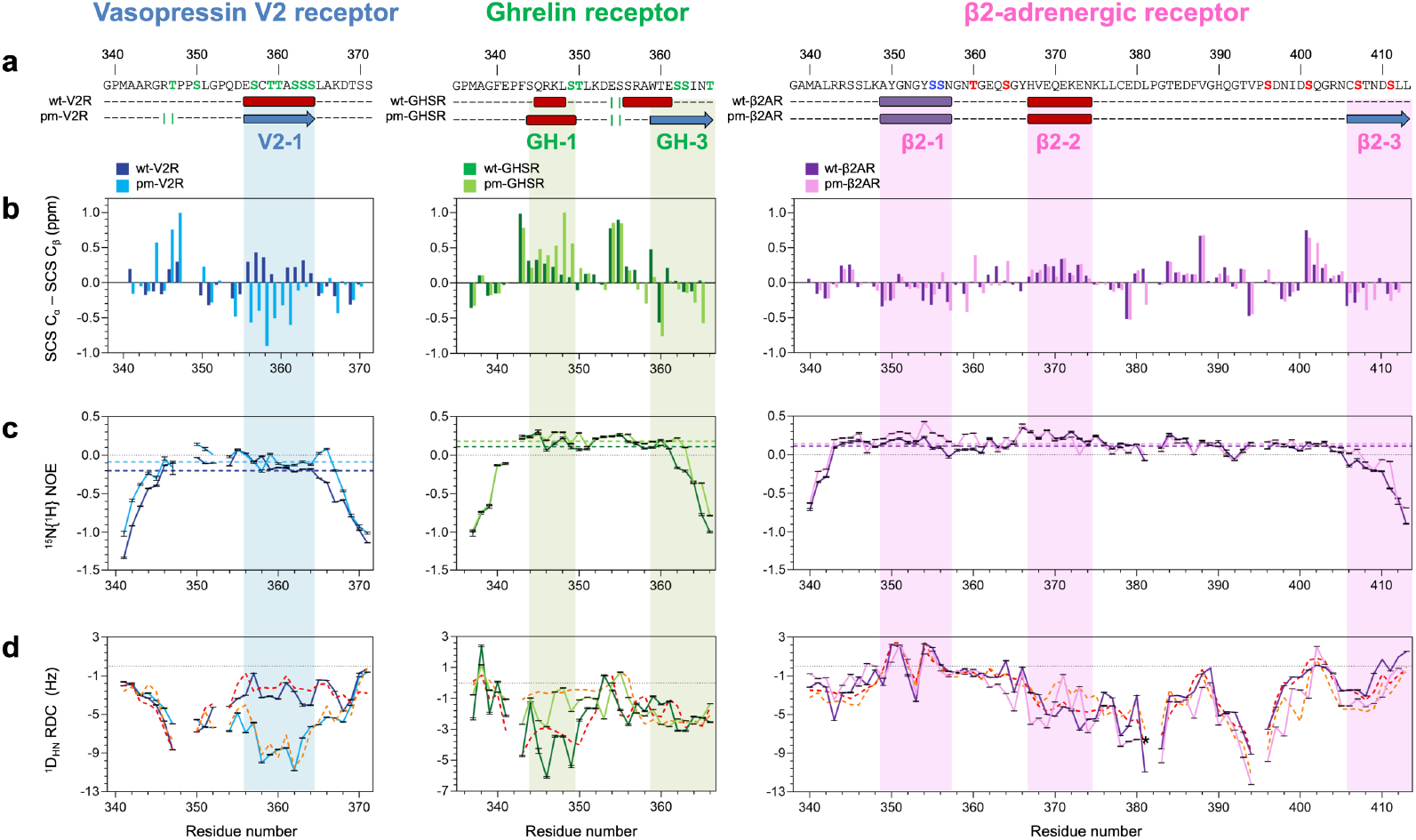
Comparison of the structural features of GPCR C-terminal domains in their non-phosphorylated (wt, dark color, data from (Guillien et al., 2022)) and phosphomimetic form (pm, light color) of V2R-Cter (blue, left); GHSR-Cter (green, middle) and β2AR-Cter (purple, right). **a** Schematic secondary structure consensus of all NMR data along the primary sequences. Known phosphorylation sites mutated to glutamic acid (E) in the variants (pm) are indicated in green, red and blue on the sequence (according to Fig. 1). Helix, β-strand, extended conformation (β-strand or polyproline helix 2, PPII) and turns are represented as red cylinder, blue arrow, purple box and green bars, respectively. **b** Computed secondary chemical shift SCS Cα – SCS Cβ. **c** ^15^N{^1^H} heteronuclear NOEs profiles from the ratio of intensities measured in saturated (I) and unsaturated (I_0_) spectra. Averages are in dash colored line. **d** Experimental and back-calculated ^1^D_NH_ RDC. Comparison of experimental RDCs measured in bacteriophage Pf1 for GHSR-Cter, alcohol mixtures for V2R-Cter and β2AR-Cter. Back-calculated RDCs from biased ensembles generated with FM (dashed lines) for wt (red) and pm (orange). The peak displaying signal overlap was removed (dark star).

### pm-GHSR-Cter

The SCS profile of GHSR-Cter phosphomimetic (pm) variant revealed the presence of an α-helix (from residues 345 to 348 - called GH-1) and a β-strand conformation (from 359 to 366 - called GH-3) with positive and negative SCS values, respectively (Fig. 2b). Previously, we identified two helices for the non-phosphorylated (wt) form; i.e. GH-1 (from residues 345 to 348) and GH-2 (from residues 356 to 361) (Fig. 2a) (Guillien et al., 2022). SCS values of the pm variant in GH-1 region were higher than in the wt peptide, suggesting a stronger helical structuration. Interestingly, GH-2 helix found in the wt peptide was shifted downstream and converted to a β-strand conformation (GH-3) (Fig. 2b). The transition from a helical to a β-strand conformation was confirmed by the decreased of ^3^JHNHA scalar couplings with less than 6 Hz (from 46% to 29%) and a small increase of couplings larger than 8 Hz (from 3% to 4%) between the wt and the pm forms (Fig. S3a). Also, scalar couplings in GH-3 were higher for the pm variant than for the wt peptide, which is in line with a tendency to form a β-strand conformation (Fig. S3b). Dynamic parameters showed an increase of rigidity in GH-1 and GH-3 regions for the pm variant, with higher averaged heteronuclear NOE (0.18 ± 0.01) and transversal relaxation R2 rate (3.33 ± 0.04 Hz) compared to the wt (0.11 ± 0.01 and 3.17 ± 0.06 Hz, respectively) (Figs. 2c and S4). Conformational changes between the wt and pm peptides were also highlighted by RDC measurements. In agreement with SCS data, the comparison of RDC profiles showed an increase of RDCs in GH-1 and a slight decrease of these in GH-3 (Fig. 2d), suggesting a stronger helical structuration in GH-1, and a switch and shift from a helical (wt) in GH-2 to a β-strand conformation (pm) in GH-3.

### pm-β2AR-Cter

The SCS profile of the β2AR-Cter pm variant highlighted three regions with negative and positive values indicating transient secondary structures: an extended conformation from residues 349 to 357 (called β2-1), a helix from 368 to 376 (called β2-2) and β-strand conformation from 407 to 413 (called β2-3) (Figs. 2 and S3-S5). The NMR characterization of the wt peptide gave similar results for the two first regions (Guillien et al., 2022) while no secondary structure was observed in the third identified region (Fig. 2B). The appearance of a β-strand in β2-3 was manifested by the increase of ^3^JHNHA values between 7 and 8 Hz (from 13 to 28 %) (Fig. S3). Moreover, in this region, heteronuclear NOE and transversal relaxation R2 data showed a slight increase of their values in the pm variant compared to the wt form, while the longitudinal relaxation R1 remained unchanged (Figs. 2C and S4). These results suggested a local structuration and reduced conformational mobility in agreement with the presence of a transient secondary structure in β2-3. The emergence of β2-3 was also observed in the RDC analysis, RDCs adopted more negative values in β2-3 for the pm variant compared to the wild type, suggesting the presence of a β-strand conformation (Figs. 2d and S5). Altogether, comparison of all β2AR-Cter’s NMR data in their non-phosphorylated (wt) and phosphomimetic (pm) forms confirmed a conformational transition of the β2-3 region from a random coil (non-phosphorylated form) to a β-strand (phosphomimetic form) (Fig. 2a). Finally, no drastic changes were observed in the long-range interaction previously identified in wt-β2AR-Cter (Fig. S6).

### Conformational ensemble of GPCR-Cters phosphomimetic forms

To go further in the description of pm-GPCR-Cter conformations, we generated random-coil ensembles of 50,000 conformers for each pm variant using Flexible Meccano (FM) (Bernadó et al., 2005; Ozenne et al., 2012a). The quality of these ensembles was evaluated by comparing back-calculated to experimental RDCs. Back-calculated RDCs from random-coil ensemble showed discrepancies with the experimental ones for all pm variants, especially in beta-stranded regions V2-1, GH-3 and β2-3 (Fig. S5b). Thus, we included conformational preferences defined by NMR data in biased ensembles of 50,000 conformers (Figs. 2d and S5b). The best agreement for pm-V2R-Cter ensemble (χ^2^ = 1.23) was found when imposing: 10%, 50% and 25% of β-strands for residues 352-357, 359-362 (V2-1) and 364-368, respectively, (Fig. S5b), while wt-V2R-Cter ensemble revealed a 5% helix content in V2-1. Thus, ensemble description of V2R-Cter clearly confirmed the conformational change between pm- and wt-V2R-Cter and a strong propensity of the pm variant to adopt a β-strand conformation in V2-1. In addition, the two poly-proline II helices (PPII) identified in the non-phosphorylated (wt) ensemble were also added in position 344-345 (20%) and 350-351 (30%) of the pm-variant. In the best ensemble of pm-GHSR-Cter (χ^2^ = 0.77), the major difference between the pm variant and the wild-type ensembles was the introduction of 60% of β-strand in GH-3 (Fig. S5b) instead of 8% of helix in GH-2, respectively. It confirmed the structural transition from a helix (wt) to a β-strand (pm) in GHSR-Cter. In addition, a stronger helical structuration was observed in GH-1 variant (4% of helix from 349 to 351) (Fig. S5b), compared to wild-type ensemble, where no conformational bias was required in this region. For the β2AR-Cter variant, the best ensemble (χ^2^ = 4.21) was obtained by imposing 5% of β-strand near β2-3 region (Fig. S5b), while no conformational constraints were added in this region for the wild type. This result showed that the conformational change to a β-strand also occurred in β2AR-Cter variant but to a lower extent than in the two other Cters. Another difference appeared in β2-2 region where 5% helix was added instead of a PPII in the wild type. The ensemble of the pm variant was also constrained by a 10% β-turn I in residues 401-402 (Fig. S5b). For both pm- and wt-β2AR-Cter, a long-range contact (15 Å) was incorporated to the model. Indeed, decreases in PRE ratio were identified between regions surrounding β2-1 (residues 338-357) and β2-2 (residues 367-386) (Fig. S5c) for both pm- and wt-β2AR-Cter. Altogether, wild-type ensembles show a conformational transition occurring when introducing phosphomimetic residues involving larger populations of β-strand conformations.

### Interaction of the C-terminal domain of GPCRs with Arrestin

To assess the functional impact of the conformational changes described above, we then analyzed the interaction of the three GPCR-Cter with their major biological partner, arrestin.

#### Arrestin binding sites contain putative phosphorylation motifs

NMR is a suitable tool to study the location and strength of transient interactions by monitoring the changes in chemical shift and/or intensity of a labeled protein with increasing amounts of the unlabeled partner (Schumann et al., 2007; Williamson, 2013). With this aim, we recorded a set of ^15^N-HSQC spectra of ^15^N labeled GPCR-Cters incubated with different ratios of unlabeled arrestin-2 to perform a Chemical Shift Perturbation (CSP) assay. For the three phosphomimetic variants, changes in chemical shifts were observed when comparing the spectra of the protein in the absence and presence of arrestin-2 (respectively 1:0 and 1:10) (Fig. S6). For the pm-V2R-Cter, chemical shift changes and an intensity decrease proportional to the amount of added arrestin-2 were observed from residues 356 to 366 (Figs. 3 and S7a). Similar results were obtained for pm-GHSR-Cter from residues 358 to 366 (Figs. 3 and S7b). Concerning pm-β2AR-Cter, small chemical shift changes were observed for residues 408-413 at a 1:10 ratio (Figs. 3 and S7c). Importantly, no interaction was observed between the three non-phosphorylated C-terminal domains (wt-GPCR-Cter) and arrestin-2, highlighting the importance of phosphorylation for GPCR recognition of arrestin (Figs. 3 and S6-S7).

**Fig. 3.**
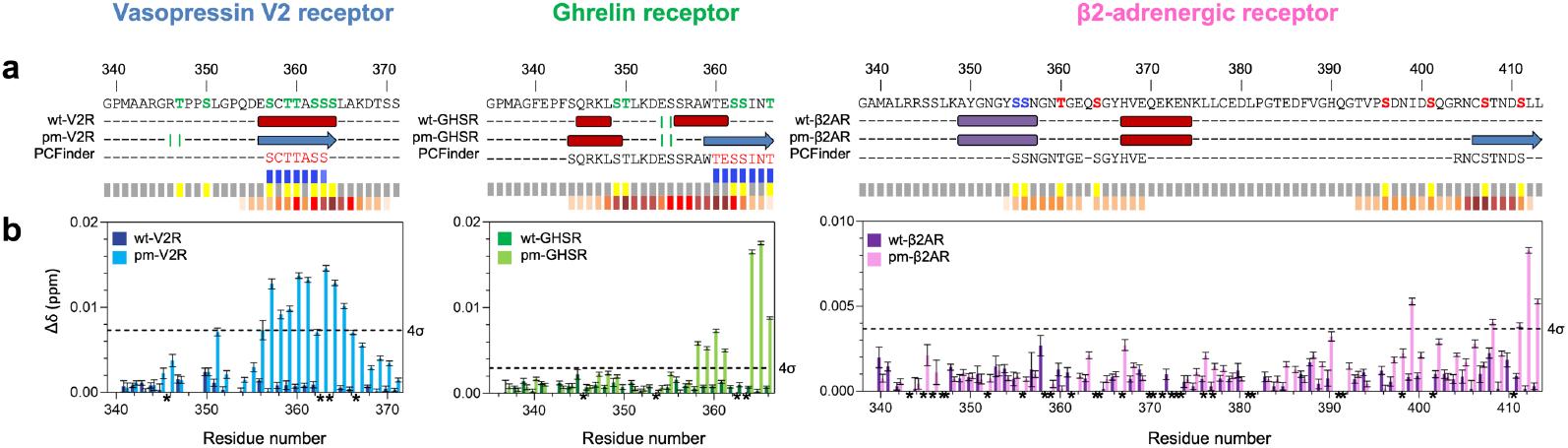
Interaction between arrestin-2 and the C-terminal domains of V2R, GHSR and β2AR in their non-phosphorylated (wt, dark colors) and phosphomimetic (pm, light colors) forms. For each receptor: **a** sequence of C-terminal domain is indicated according to Fig. 1 and just below successively, secondary structure contents of non-phosphorylated (wt) and phosphomimetic (pm) variants, and predicted phosphorylation motifs (PCFinder = PhosCoFinder (X Edward Zhou et al., 2017)); (in the middle) putative phosphorylation motifs (p[S/T](X)Xp[S/T]XXp[S/T]/E/D motifs) displayed on three different tracks: blue track contain the complete motifs (motif containing three negatively charged residues), grey track contain the sequence with the known phosphorylation sites from Uniprot and PhosphoSitePlus database (Hornbeck et al., 2015) in yellow, reddish track contain partial motifs (motif containing two negatively charges residues instead of three). For partial motifs, darker colors indicate residues involved in multiple motifs. **b** ^1^H/^15^N Chemical Shift Pertubation (CSP) observed at ratio 1:10 (10 μM GPCR-Cter:100 μM arrestin) aligned with the putative phosphorylation motifs. CSP were recorded on sample at 10 μM with 0 or 100 μM of arrestin-2 at 800 MHz and 20 °C in a 50 mM Bis-Tris pH 6.7, 50 mM NaCl buffer. Cut-offs are in black dashed lines and peaks displaying signal overlap were removed (dark star).

Noteworthy, the identified binding regions overlapped with the regions undergoing a conformational transition between the non-phosphorylated (wt) and the phosphomimetic (pm) variants (Fig. 3). Moreover, these regions were also predicted to contain phosphorylation motifs using the server PhoCoFinder (X Edward Zhou et al., 2017). These phosphorylation motifs were proposed as motifs required for high-affinity arrestin binding. For pm-V2R-Cter and pm-GHSR-Cter, the binding region encompassed a so-called complete phosphorylation motif (three negatively charged residues), while pm-β2AR-Cter contained only a partial phosphorylation motif (two negatively charges residues instead of three) (Fig. 3). This was in line with the short binding region with a weaker chemical shift perturbation (0.009 ppm ± 0.001), suggesting a lower affinity for arrestin-2. On the opposite, the phosphomimetic variant of V2R-Cter experienced the largest chemical shift perturbation with a Δδ of 0.015 ppm (± 0.001) and the largest binding region with a dozen of impacted residues, which should be related to a stronger interaction with arrestin-2. These observations provide the structural bases of the previously reported higher affinity for arrestin of V2R with respect to GHSR1a and β2AR (Oakley et al., 2000b).

#### GRK phosphorylation acts as a switch of GPCR’s C-terminal domains conformation

We thus showed that the regions that exhibited conformational transition from helical or random coil to β-strand were involved in the interaction with arrestin. Two different mechanisms, conformational selection or induced fit, can then be considered. Indeed, the recognition with arrestin could occur due to a pre-configuration of the C-terminal domain of GPCRs upon GRK phosphorylation or, alternatively, the final configuration could be induced by the folding of the C-terminal domain of GPCRs upon binding to arrestin. To discriminate between both processes, we checked if there was a change in conformation before and after binding of arrestin. For the phosphomimetic variant of V2R-Cter, secondary chemical shift (SCS) profiles in presence and in absence of arrestin were similar (Fig. 4), suggesting that the interaction did not induce a folding, pleading for a conformational selection mechanism. This hypothesis was supported by the observation that the non-phosphorylated forms, which did not encompass either any charges nor β-strand conformation, did not interact with arrestin, and by the X-ray structures of GPCR:arrestin complexes where the bound state of the Cters displayed a β-strand conformation. In our suggested model, a pre-existing with well-disposed charges along the C-terminal domain of GPCRs would be necessary for arrestin recognition.

**Fig. 4.**
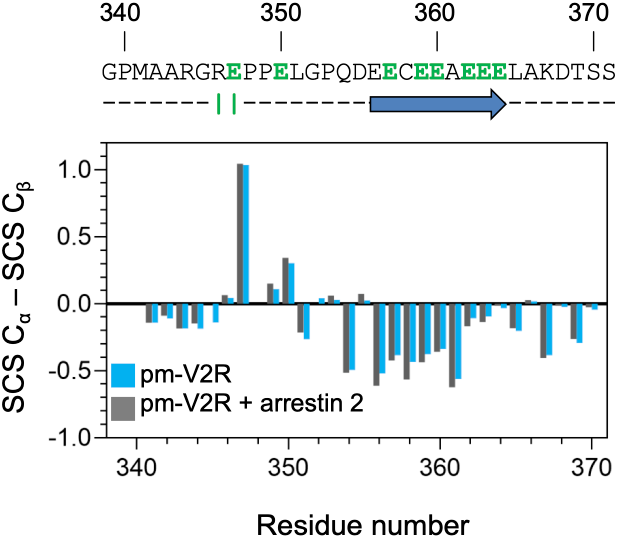
Conformational selection of phosphomimetic V2R-Cter by arrestin-2. ^13^Cα/^13^Cβ secondary chemical shift (SCS) computed with POTENCI (Nielsen and Mulder, 2018; Tamiola et al., 2010) of pm-V2R-Cter in absence (blue) or in presence of arrestin (grey). 3D spectra were recorded on ^13^C/^15^N pm-V2R-Cter (50 μM) in absence or in presence of arrestin-2 (250 μM) at a ratio of 1:5 at 800 MHz and 20 °C in a 50 mM Bis-Tris pH 6.7, 50 mM NaCl buffer.

#### Arrestin reorganization in GPCR-Cter:arrestin complexes

We then analyzed if, conversely, interaction with the pm peptides affected the conformational features of arrestin. To this end, we first developed an arrestin-activation sensor based on the FRET signal between a fluorescence donor (AlexaFluor350) and an acceptor (AlexaFluor488) attached to each lobe of purified arrestin-2 (Fig. 5a). The selected positions, *i*.*e*. V167 and L191, are closely related to those recently shown to report on arrestin activation, based on single molecule fluorescence experiments (Han et al., 2021). A significant change in the FRET ratio was observed (Fig. 5b) in the presence of the phosphomimetic peptides, suggesting that the latter interact with purified arrestin. In contrast, no significant change in the FRET ratio was observed for the non-phosphorylated peptides, suggesting that these peptides do not interact with arrestin. However, an interaction that would not be accompanied by a change in the conformation of arrestin cannot be excluded at this stage. To be noted, for the GHSR- and β2AR-Cters, a statistically significant FRET ratio change was observed only at high peptide-to-arrestin molar ratios, suggesting a significantly lower affinity of these peptides compared to the V2R-Cter. However, in all cases, the high excess required to get significant FRET ratio differences suggested a weak interaction between arrestin and the phosphomimetic peptides. Indeed, the affinities (KD) in these transient complexes were estimated in the ∼ 200 μM range using microscale thermophoresis (MST) and fluorescence anisotropy (Fig. S8). For instance, pm-GHSR-Cter, which showed a clear interaction with arrestin-2 by NMR (Figs. 3 and S7b), could not be saturated in MST or fluorescence anisotropy experiments (Fig. S8).

**Fig. 5.**
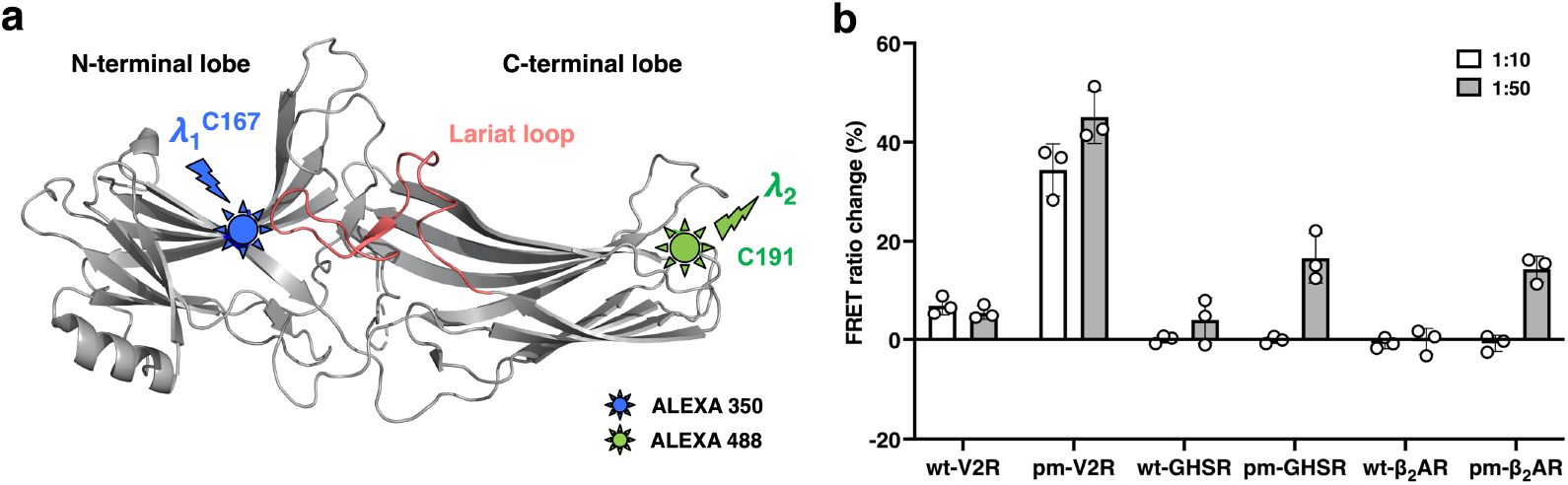
FRET-monitored interaction between purified arrestin-2 and the non-phosphorylated and phosphorylated Cters. **a** Structure of arrestin2 (PDB ID: 1G4M) with fluorescence donor (AlexaFluor350) and an acceptor (AlexaFluor488) attached to each lobe. **b** Changes in the FRET ratio measured for the dual labeled arrestin in the absence and presence of the wt peptides (wt-V2R-Cter, wt-GHSR-Cter, wt-β2AR-Cter) and their phosphomimetic counterparts (pm-V2R-Cter, pm-GHSR-Cter, pm-β2AR-Cter) at different arrestin-to-peptide stoichiometric ratios (1:10 or 1:50). Data expressed in % are the mean ± SD of three experiments.

### Phosphomimetic peptides mimic the interactions made by phosphopeptides

We finally performed molecular dynamics (MD) simulations to understand the interactions between the phosphomimetic peptides and arrestin-2. We used the NMR data to define the regions of peptide that bind to arrestin-2 and generated the initial conformations of only the binding region of peptide in complex with arrestin-2 using the structure of V2R phosphorylated peptide (V2Rpp) bound to arrestin-2 (PDB ID: 4JQI) (Shukla et al., 2013a). These conformations were refined using FlexPepDock server which led to the formation of a β-sheet at the binding regions of both pm-V2R-Cter and pm-GHSR-Cter while pm-β2AR-Cter adopted a less clear β-sheet (Fig. 6). This agrees with the secondary structure determination from the NMR data (Fig. 2 and Fig. S5b). The refinement by FlexPepDock was followed by the addition of the rest of the residues of the peptides and performing MD simulations by gradually releasing the restraints from the complex (see Methods). For both pm-V2R-Cter and pm-GHSR-Cter, the β-strand at the primary binding regions was consistently maintained and the interactions with arrestin-2 via an anti-parallel β-sheet were preserved throughout the simulations (Fig. 6a-b). For pm-β2AR-Cter, the binding region was more flexible and showed occasional detachment and rebinding with more than 90% frames in the bound state. On the other hand, the rest of the three peptides showed much more conformational flexibility and made transient interactions with multiple regions of arrestin-2 (Fig. 6a). The probability of contacts between residues of phosphomimetic peptides and arrestin showed that peptides make transient interactions with all the pockets identified on the surface of arrestin (Figs. 7 and S9a). We observed the formation of almost all the hydrogen bonds (H-bonds) present in the crystal structure of V2Rpp (Shukla et al., 2013a) including H-bonds with K10-11, R25, K107, K146, K160, R165 and K294 (Fig. 6c).

**Fig. 6.**
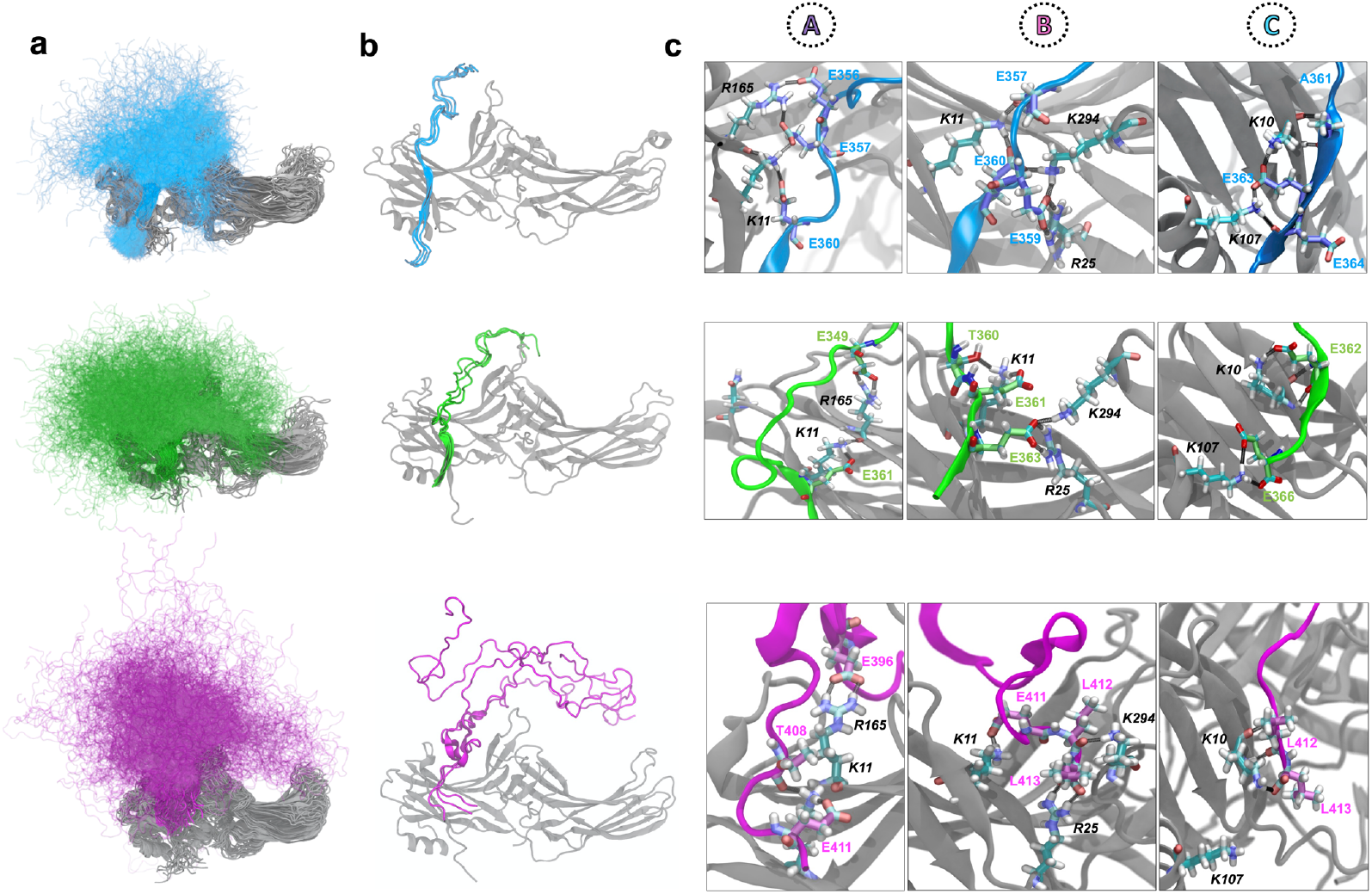
Modeling of pm-V2R-Cter (blue), pm-GHSR-Cter (green) and pm-β2AR-Cters (pink): arrestin (grey) complexes. **a** The conformational ensemble obtained from MD simulations for the three phosphomimetic Cter peptides in complex with arrestin. Every 500^th^ conformation from the four independent simulations is shown. **b** Three selected peptide conformations from the calculated ensembles which form the highest number of hydrogen bonds with arrestin. **c** The insets show the different hydrogen bonds formed by the phosphomimetic Cters with arrestin pockets. Arrestin pockets (A, B and C) are described in Fig. 7. Residues from arrestin are labeled in black and residues from the peptides are labeled in their respective colors.

**Fig. 7.**
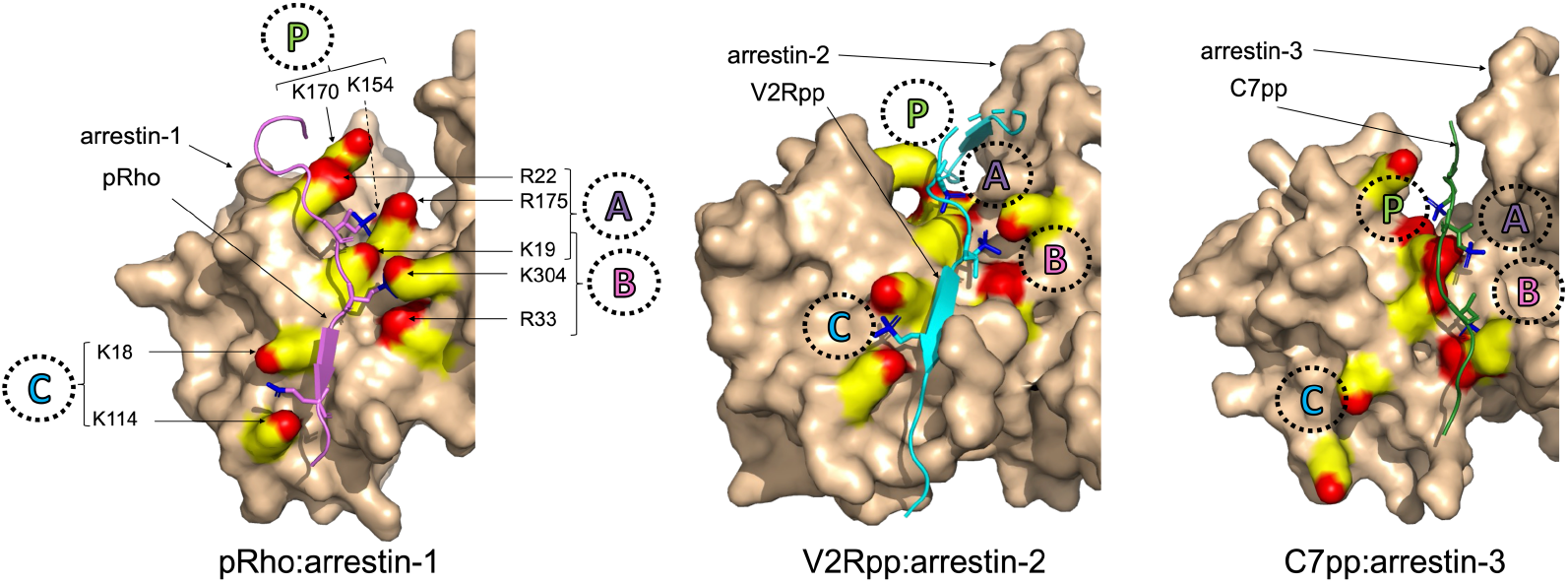
Atomic structures of GPCR:arrestin complex. For the rhodopsin:arrestin-1 complex (X Edward Zhou et al., 2017), the C-terminal domain was phosphorylated during its production in insect cells (pRho); while in the two other complexes, V2R and CXCR7, the C-terminal domain correspond to synthetic fully phosphorylated peptides (respectively V2Rpp and C7pp) (Min et al., 2020; Shukla et al., 2013a). V2Rpp and pRho interact with arrestin pocket A (K19, R22 and R175), B (K19, R33 and K304) and C (K18 and K114) while C7pp interacts with pocket P (K154 and K170), in addition of pocket A and B. These pockets are conserved among arrestins and residue numbers correspond to the sequence of human arrestin-1. K154 is localized in a crest behind R175 and is not visible in the figure (dashed arrow).

This demonstrates that the phosphomimetic peptides can recapitulate the interactions made by phosphopeptides and are likely to activate arrestin-2 by the same mechanism (Fig. S10a). Moreover, truncated C-terminal domain of V2R adopt a similar conformation when it is a continuation of its receptor’s amino acid sequence (Bous et al., 2022), which could support the generality of the proposed mechanism of arrestin binding for Rhodopsin like receptors (Fig. S10b). We also observed that the total number of H-bonds between the phosphomimetic peptide and arrestin is the highest for pm-V2R-Cter followed by pm-GHSR-Cter and the lowest for pm-β2AR-Cter, which agrees with the experimentally observed affinity trend (Fig. S9b). Interestingly, some of the more subtle features such as the bifurcated salt-bridge observed for K294 with derived V2Rpp (Baidya et al., 2020), which has been shown to be essential for keeping the lariat loop (Fig. 5a) in the active conformation, was also reproduced by pm-V2R-Cter. This bifurcated salt-bridge was formed by E357 and E360 with an additional H-bond with C358 (Fig. S9c).

## Discussion

Understanding how GRK phosphorylation of the C-terminal domain of GPCRs (GPCR-Cter) dictates the recruitment of arrestin is pharmacologically and clinically important. The phosphorylation-barcode model was proposed a decade ago, stating that the phosphorylation pattern on receptor’s C-terminal domain directs arrestin binding and subsequent functions (Nobles et al., 2011b). However, how phosphorylation of GPCR-Cter at specific sites leads to distinct arrestin conformations and signaling outcomes remains an open question. Here, we studied how GRK phosphorylation affects the conformation of GPCR C-terminal domains and their interaction with arrestin-2 using a phosphomimetic strategy (Pearlman et al., 2011). Using a divide and conquer strategy, we have unraveled their complex mechanism of action. For this, we used C-terminal constructs of the receptor to be able to extract high resolution information of these extremely flexible systems (Xie et al., 2007). Thus, C-terminal domains of GPCRs are not visible in full-length receptors free in solution due to technical limitation of the structural biology tools used so far. The strategy of divide and conquer has been previously used in GPCR studies such as the V2pp:arrestin complex (He et al., 2021b; Shukla et al., 2013a), as well as for other macromolecular complexes (Cordeiro et al., 2019; Senicourt et al., 2021). Furthermore, we used phosphomimetics in order to work with a homogeneous system, facilitating their structural characterization (Bibow et al., 2011).

Of importance, our data show that the phosphomimetic C-terminal domain variants are able to interact with arrestin-2, which is not the case of the unmodified ones. Besides being in line with the fact that phosphorylation is generally necessary for arrestin binding (Gurevich and Gurevich, 2019), this indicates that the phosphomimetic models are functional and can therefore be used as a tool to study GPCR:arrestin complexes and their associated arrestin-dependent signaling pathways. Indeed, the modeling of pm-GPCR-Cter:arrestin complexes and MD simulations show that the binding mode of the phosphomimetic V2R C-terminal domain to arrestin is similar to phosphopeptides (Shukla et al., 2013a), or to the whole V2 receptor in complex with arrestin (Bous et al., 2022), as they are able to make all the interactions with different regions of arrestin. This further demonstrates the suitability of isolated C-terminal phosphomimetic peptides as models for studying GPCR-arrestin interactions as they exhibit a similar mode of arrestin activation.

Our data show that both the non-phosphorylated C-terminal domains and their phosphomimetic counterparts behave in solution as intrinsically disordered regions (IDRs) with transient secondary structure elements. However, mimicking GRK phosphorylation causes the formation of β-strand conformations in localized segments of the GPCR-Cters, suggesting that the GPCR-Cter functional folding is regulated by GRK phosphorylation, similar to other proteins such as the eukaryotic translation factor 4E-binding protein 2 (Bah et al., 2015). Noteworthy, the β-strand conformations in the three C-terminal domains, only present in the phosphomimetic variants, corresponded to the arrestin interacting region (Fig. 3). Interestingly, several atomic resolution structures, obtained by X-ray crystallography and cryo-electron microscopy, showed a β-strand conformation of the fully phosphorylated C-terminus peptide of vasopressin V2 receptor (V2pp), which formed an anti-parallel β-sheet with the N-domain of arrestin-2 (He et al., 2021a; Nguyen et al., 2019; Shukla et al., 2013b; Staus et al., 2020). Until now, only structures of class B receptors, which interact more strongly with arrestin, or chimeric class A receptors with the C-terminus of V2R (V2Rpp), are reported in the literature. Here, we provide two models of class A receptor that interact transiently with arrestin.

A similar fold was also observed for the phosphorylated rhodopsin C-terminus (pRho) in interaction with visual arrestin-1 (X Edward Zhou et al., 2017). In these complexes, an extensive network of electrostatic interactions exists between the negatively charged residues of the C-termini and three conserved pockets of positive residues in the N-domain of arrestin: A (19, R22 and R175), B (K19, R33 and K304) and C (K18 and K114) (Fig. 7; residue number corresponding to the human arrestin-1). V2Rpp and pRho β-strands are located between arrestin pockets B and C. However, in a recent X-ray atomic structure showing a synthetic fully phosphorylated peptide of CXCR7 receptor (C7pp) in interaction with arrestin-3 (Min et al., 2020), the C-terminal domain of CXCR7 adopted an elongated loop that, in addition to pockets A and B, interacts with a new pocket P (K154 and K170, Fig. 7). Moreover, Yu and collaborators showed that the interaction of phosphate of V2R C-terminus with specific pocket of arrestin affects the interaction surfaces of arrestin with its cytosolic partners (He et al., 2021c). Thus, it is possible that, depending on the conformation of the GPCR C-terminal domain, phosphates do not interact with the same pockets of arrestin, leading to distinct arrestin function. This folding could also be used to select a particular arrestin subtype.

Binding regions identified in our study contained phosphorylation motifs specific to GPCRs (PhosCoFinder (X Edward Zhou et al., 2017) and Fig. 3). These phosphorylation motifs correspond to a long p[S/T]XXp[S/T]XXp[S/T]/E/D motif or short p[S/T]Xp[S/T]XXp[S/T]/E/D motif. It was proposed that the presence of three negatively charged residues (full motifs) increased arrestin interaction (X Edward Zhou et al., 2017). Conversely, the absence of one of the three charges (partial motif) reduces the interaction. The vasopressin V2 and the ghrelin receptors encompass full motifs, while the β2-adrenergic receptor displayed only two clusters of partial motifs (Fig. 3). This was consistent with our NMR observations where V2R-Cter encompassed the largest binding region, whereas β2AR-Cter presented the shortest binding region and the lowest arrestin binding affinity. Low affinities were previously observed between phospho-peptides of V2R-Cter and arrestin (He et al., 2021b) and between β2AR receptor and arrestin (Gurevich et al., 1995; Min et al., 2020). In addition, it has been already shown in IDPs that multisite phosphorylation modulates their affinity toward partners (Van Roey et al., 2012). For example, successive phosphorylation events of the p53 transcriptional factor increase its recruitment toward the CREB-binding protein (Lee et al., 2010). Thus, it has been proposed that multisite phosphorylation acts as a rheostat to enhance binding of IDRs/IDPs to their partner. A similar binding mode could occur between GPCR-Cters and arrestin where multivalency of phosphates would enhance the affinity of GPCR to arrestin and stabilize the interaction. This would be in agreement with the p[S/T](X)Xp[S/T]X(X)p[S/T]/E/D motif where it has been proposed that three phosphates are required to have higher affinity binding to arrestin (X Edward Zhou et al., 2017). This would also be in agreement with the work of Yu and coll. showing that the absence of only one phosphate reduces the affinity of V2R to arrestin (He et al., 2021b).

Our study highlights a conformational transition between the basal non-phosphorylated state and the phosphomimetic state from a helical or random coil to a β-strand conformation. This transition occurs in the region that interacts with arrestin-2. Moreover, no additional conformational changes were observed in the bound state of the variant of V2R-Cter, compared to its free state, suggesting conformational selection as the predominant interaction mechanism.

Based on this evidence, a model emerges where GRK phosphorylation dictates the final configuration of GPCR C-terminal domains to trigger the binding mode to arrestin. The selected arrestin conformation resulting from the interaction of GPCR phosphate with specific pockets of arrestin would expose specific interaction surfaces recognized by different cytosolic partners, and thus, modulate arrestin function. Hence, in this model, the signaling fate in the arrestin pathway would, at least in part, also depend on the structural impact of GRK phosphorylation on its partners.

## Materials and Methods

### Expression and purification

#### Recombinant human GPCR-Cter

Phosphomimetic GPCR C-termini genes were purchased from GeneArt gene synthesis (Life technologies). They were subcloned into pETM33 vector for GHSR-Cter and V2R-Cter and PET1a vector for β2AR-Cter. The constructs were fused with a (His)6-GST-tagged followed by the HRV 3C protease recognition site (for GHSR-Cter and V2R-Cter) or by the TEV protease recognition site (for β2AR-Cter). The three GPCR C-termini were expressed and purified as described previously (Guillien et al., 2022).

#### Recombinant rat arrestin-2

pETM11-arrestin-2 plasmid was transfected into *E. Coli* BL21 DE3 strain (ThermoFisher Scientific). The cells were cultured and induced with 0.25 mM isopropyl-β-D-thiogalacoside (IPTG) at a OD600 of 0.5 nm. After growing over-night at 20 °C, bacteria were harvested, resuspended and lysed by French-Press. Cell debris was removed by centrifugation and the soluble fraction was loaded onto 5 mL HisTrap HP column (Cytiva) in buffer A (50 mM Tris pH 8, 200 mM NaCl and 2 mM DTT). The protein was recovered using step elution with buffer A supplemented with 500 mM imidazole. The eluted protein was dialyzed over night at 4 °C in buffer A with TEV protease (ratio protease:protein 1:50 (w/w)). Protease and His-TEV tag were removed using HisTrap HP column. Untagged proteins were diluted to 50 mM NaCl and injected into HiTrap Q HP (Cytiva) in buffer C (20 mM Tris pH 8, 50 mM NaCl and 2 mM DTT). The protein was eluted with a linear gradient of buffer C supplemented with 1M NaCl. Proteins were concentrated using a 10 kDa Millipore concentrator and injected into a HiLoad 16/60 Superdex 200 size exclusion column (Cytiva) in 50 mM Bis-Tris pH 6.7, 50 mM NaCl and 2 mM DTT. The purified proteins were then concentrated using Millipore concentrator and aliquots were flash-frozen in 50 mM Bis-Tris pH 6.7, 50 mM NaCl, 1 mM EDTA (EthyleneDiamineTetraAcetic acid), 0.5 mM TCEP (Tris(2-CarboxyEthyl)Phosphine) and stored at -80 °C.

#### Recombinant arrestin-2 for FRET experiments

The full-length cysteine-free arrestin-2 mutant was modified with two cysteines at positions 167 and 191 and an N-terminal hexahistidine tag followed by a 3C protease site. The sequence was codon-optimized for expression in *E. coli* and cloned into a pET-15b vector (Genecust). NiCo21(DE3) competent *E. coli* (NEB) cells were transformed with the resulting expression vector. Cultures were grown in 2YT medium supplemented with 100 μg/mL carbenicillin at 37°C up to an OD600 of 1.0. The temperature of the culture was then decreased to 16°C and protein synthesis induced with 25 μM IPTG for 20 hours. Cells were harvested and resuspended in ice-cold lysis buffer (50 mM Tris-HCl pH 8, 300 mM NaCl, 15% glycerol, 1 mM TCEP) supplemented with a protease inhibitor cocktail (Roche). Cells were lysed by sonication, centrifuged at 27,000xg for 45 minutes and the supernatant was applied to a HisTrap 5 mL column (GE Healtcare). The resin was washed with a 50 mM Tris-HCl pH 8, 300 mM NaCl, 15% glycerol, 1 mM TCEP buffer, with the same buffer supplemented with 20 mM imidazole then with the same buffer containing 40 mM imidazole. The protein was eluted with the same buffer containing 200 mM imidazole and dialyzed into a 25 mM Na-HEPES, 200 mM NaCl, 1 mM TCEP, 10% glycerol, pH 7.5 buffer. The protein after dialysis was digested with 3C protease at a protease:arrestin ratio of 1:10 (w:w) for 16 hours at 20 °C and subjected to reverse-nickel purification. The protein in the flow-through fractions were recovered, diluted with the same buffer devoid of NaCl to reach a final salt concentration of 40 mM and loaded on a HiSure Q 5 mL column (GE Healthcare). After washing with a 25 mM Na-HEPES, 40 mM NaCl, 1 mM TCEP, 10% glycerol, pH 7.5 buffer, elution was carried out with a linear 40 to 400 mM NaCl gradient in 25 mM Na-HEPES, 1 mM TCEP, 10% glycerol, pH 7.5. The fractions in the elution peak were pooled, concentrated using a 30 kDa spin concentrator (Amicon) and subjected to size-exclusion chromatography using a Superdex 200 increase 10/300 GL column (GE Healthcare) in a buffer 20 mM HEPES pH 7.5, 200 mM NaCl, 10% glycerol, 1 mM TCEP. The elution peak fractions were pooled and concentrated using a 30 kDa spin concentrator (Amicon) up to a concentration of 10-20 μM. A sequential labeling strategy was then used to label arrestin with the FRET probes (Zosel et al., 2022). Purified arrestin was first incubated with 0.8 equivalents of AlexaFluor350 maleimide (ThermoFisher) for 3 hours in the dark. Unreacted fluorophore was removed on a Zeba spin desalting column (ThermoFisher). 1.3 equivalents of AlexaFluor488 maleimide (ThermoFisher) were then added and incubated for 3 additional hours in the dark. The reaction was stopped with 5 mM L-cysteine and unreacted fluorophore again removed on a Zeba spin column (ThermoFisher). The degree of labeling was determined using the known extinction coefficients of the protein and both fluorophores at their absorbance maxima. The typical ratio of donor to arrestin was in the 0.8-0.9 range, with a typical acceptor/donor labeling ratio in the 1-1.2 range.

### Biophysical characterization

#### Size Exclusion Chromatography – Multi-Angle Light Scattering (SEC-MALS)

The experiments were performed at 25 °C using a Superdex 75 10/300 GL column (Cytiva) connected to a miniDAWN-TREOS light scattering detector and an Optilab T-rEX differential refractive index detector (Wyatt Technology, Santa Barbara, CA). The column was equilibrated in 50 mM BisTris pH 6.7, 50 mM NaCl, 1 mM TCEP and 0.5 mM EDTA buffer filtered at 0.1 μM, and the SEC-MALS system was calibrated with a sample of Bovine Serum Albumin (BSA) at 1 mg/mL. Samples were prepared at 1.16 mM, 1.30 mM and 0.35 mM for pm-V2R-Cter, pm-GHSR-Cter and pm-β2AR-Cter, respectively. For each GPCR-Cter, 40 μL of sample were injected at 0.5 mL/min. Data acquisition and analyses were performed using the ASTRA software (Wyatt).

#### Circular Dichroism (CD) spectroscopy

Far UV-spectra of the three pm-GPCR-Cters were recorded in a quartz cuvette (path length 0.1 cm) at 0.08 mg/mL in H2O at 20 °C using a Chirascan. The ellipticity was scanned from 190 to 260 nm with an increment of 0.5 nm, an integration time of 3 s, and a constant band-pass of 1 nm. Data were treated using Chirascan and, after subtraction of the buffer signal, were converted to mean residue ellipticity ([θ]MRW, mdeg.cm^2^.dmole^-1^) using equation (1) (Chemes et al., 2012):

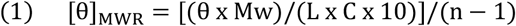

where θ is the ellipticity (mdeg), Mw is the molecular weight (g/mol), L is the cuve lengh (cm), C is the protein concentration (mg/mL) and n is the number of peptid bond.

*Small-angle X-ray scattering (SAXS) measurement and analysis*. Synchrotron radiation small-angle X-ray scattering (SAXS) data were acquired for pm-GPCR-Cters at the SWING beamline at the SOLEIL synchrotron (Saint-Aubin, France) (Thureau et al., 2021) using an X-ray wavelength of 1.03 Å and a sample-to-detector distance of 2.00 m. Samples were measured at 15 °C and at two concentrations, 5 mg/mL and 10 mg/mL, for all GPCR-Cters, in 50 mM BisTris pH 6.7, 50 mM NaCl and 2 mM DTT buffer. Before exposure to X-rays, 45 μl of sample were injected into 3 mL Superdex 75 5/150 GL column (Cytiva) at 0.2 mL/min, pre-equilibrated into the same buffer as samples. The intensity was measured as function of the magnitude of the scattering vector, s, using equation (2) (Blanchet and Svergun, 2013):

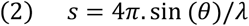

where θ the scattering angle and λ the X-ray wavelength.

The scattering patterns of the buffer were recorded before the void volume of the column (1 mL). The scattering profiles measured covered a momentum transfer range of 0.002 < *s* < 0.5 Å^−1^. Data were processed using CHROMIX from ATSAS software package to select automatically frames corresponding to buffer and sample, and perform buffer subtraction (Panjkovich and Svergun, 2018). The scaled and averaged SAXS curves were analyzed using Primus from ATSAS software package. AUTORG was used to calculate the radius of gyration.

### Nuclear Magnetic Resonance (NMR) spectroscopy

All NMR experiments were performed on Bruker Avance III 700 MHz, except 3D assignment experiments of β2AR-Cter and chemical shift perturbation (CSP) experiments at 800 MHz. The spectrometers are equipped with a cryogenic triple-resonance (^1^H, ^15^N, ^13^C) probe and shielded z-gradients. All NMR experiments were recorded, unless otherwise stated, at 20 °C in a buffer (named NMR buffer) composed of 50 mM Bis-Tris pH 6.7, 150 mM NaCl, 1 mM EDTA, 0.5 mM TCEP, 5 % D2O (Eurisotop) and 5 mM DSS-d6 (2,2-dimethyl-2-silapentane-5-sulfonate, Sigma) as internal reference (Wishart and Sykes, 1994). All experiments used the pulse sequences provided by Bruker Topspin 3.2. Squared cosine apodization was used in indirect dimensions, prior to zero-filling and Fourier transformation using TOPSPIN (version 4.0.6, Bruker) and data processing was performed using NMRFAM-SPARKY (version 1.414, (Lee et al., 2015)). For each NMR experiments, except CSP, concentrations of pm-GPCR-Cters were indicated in Table S1.

#### Backbone assignment

For the sequential backbone assignment of the ^13^C/^15^N labeled pm-V2R-Cter, pm-GHSR-Cter and pm-β2AR-Cter, HNCO, HN(CA)CO, HNCA, HN(CO)CA, CBCA(CO)NH and HNCACB triple resonance 3D experiments were recorded. HN, N, CO, Cα and Cβ nuclei of all residues were assigned, expected the first glycine N-terminal residue, the residue A339 of β2AR-Cter and proline residues.

#### Secondary Chemical Shift (SCS)

^13^Cα and ^13^Cβ chemical shifts were used to calculate SCS by subtraction of experimental chemical shifts (from the 3D experiment) from random-coil chemical shift calculated by POTENCI database (Nielsen and Mulder, 2018; Tamiola et al., 2010). To calculate SCS for interaction experiments, HNCACB triple resonance 3D experiments were recorded on a sample of 50 μM ^13^C/^15^N labeled pm-V2R-Cter free and in presence of 250 μM of arrestin-2 in a NMR buffer at 50 mM NaCl.

#### ^*3*^*JHNHA scalar coupling*

^3^JHNHA scalar coupling measurements were obtain accordingly to Vuister and Bax (Vuister and Bax, 1993). HNHA experiments were recorded in NMR buffer ^15^N-labeled pm-GPCR-Cter. Intensity of the cross-peak (Scross) and intensity of the corresponding diagonal peak (Sdiag) were extracted using Sparky. They were used to calculate the ^3^JHNHA scalar coupling of each amino acid using the equation (3):

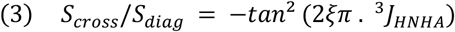

Where 2ξ is the total evolution time for the homonuclear ^3^JHNHA coupling, which has been set to 26.1 ms. ^*15*^*N NMR relaxation experiments*. Relaxation data were recorded on ^15^N-labeled pm-GPCR-Cter. Heteronuclear ^15^N{^1^H}-NOE values were determined from two experiments with on-(saturated spectrum) and off-resonance ^1^H saturation (unsaturated spectrum) which were recorded in an interleaved manner. The saturation time by 120° pulses (∼ 10 kHz) was set to 6 s and the recycle delay to 6 s. NOEs values were obtained from the ratio of intensities measured in the saturated (I) and unsaturated (I0) spectra. Longitudinal (R1) and transversal (R2) relaxation rate were measured through acquisition of ^15^N-HSQC spectra with different relaxation delays: 10, 50, 100, 200, 400, 600, 800, 1 000 ms for R1, and 16, 32, 64, 96, 160, 240, 480, 640 ms for R2. For each pick, intensity was fitted to a single exponential decay using Sparky (Lee et al., 2015) to obtain the relaxation parameters. For all relaxation parameters, three residues at N- and C-extremity were discarded from the calculation of average values due to their inherent higher flexibility.

#### ^*1*^*H-*^*15*^*N Residual Dipolar Coupling (RDC)*

Residual dipolar couplings were recorded on ^15^N-labeled pm-GPCR-Cter using 2D IPAP HSQC spectra (Ottiger et al., 1998) in isotropic and anisotropic media. For the anisotropic media, the samples were dissolved in a liquid-crystalline medium composed by 5 % (w/v) mixture of 0.85 molar ratio of polyoxyethylene 5-lauryl ether (PEG/C12E5) (Sigma) and 1-hexanol (Sigma) (Rückert and Otting, 2000) or in ∼ 20 mg/mL of filamentous phage Pf1 (Asla biotech) (Hansen et al., 1998). For pm-V2R-Cter and pm-β2AR-Cter (Fig. S5b), we analyzed RDCs in alcohol mixture (quadrupolar splitting of 33 Hz and 36 Hz, respectively), for pm-GHSR-Cter, in Pf1 medium with a quadrupolar splitting of 15.2 Hz. For more details see previous Material and Methods (Guillien et al., 2022). The ^**1**^*D*_NH_ coupling constants were obtained from the relative peak positions in the ^15^N dimension measured in the *sum* and *diff* subspectra (upfield and downfield components of the doublet). *Paramagnetic Relaxation Enhancement (PRE)*. The C378A or C406A variants of ^15^N-pm-β2AR-Cter were labeled on the remaining cysteine using 3-(2-Iodoacetamido)-proxyl (Merck). Paramagnetic samples were recorded with a recycling delay of 2 s. Reference diamagnetic samples were recorded in the same condition after addition of 5 mM fresh ascorbic acid, pH 6.7, in the NMR tube. PRE were analyzed by measuring the peak intensity ratios (Ipara/Idia) between two ^15^N-HSQC spectra of paramagnetic and diamagnetic samples, and the theoretical profile was calculated as in a strictly random coil polymer (Teilum et al., 2002).

#### Chemical Shift Perturbation (CSP)

The interaction between ^15^N-labeled GPCR C-termini and the unlabeled arrestin-2 was measured in NMR buffer containing 50 mM NaCl in order to limit the screening of potential electrostatic interactions. ^15^N-HSQC spectra were recorded on the 800 MHz spectrometer at 1:0, 1:1, 1:2, 1:5 and 1:10 ^15^N-GPCR-Cter:arrestin-2 ratio. At each point of the titration and before to record the spectrum, the complex is incubated 30 min at room temperature. For each peak in ^15^N-HSQC spectra, the intensity ratio (I/IREF) is calculated between the bound (I) and free states (IREF). Means of intensity ratio subtracted with the mean of uncertainty value was chosen as cut-off values, by excluding residues in interaction. Chemical shift perturbation Δδ were calculated using equation (4) (Grzesiek et al., 1996):

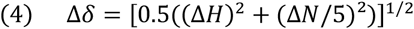

Where Δδ is the chemical shift perturbation (in ppm), ΔH is the subtraction of ^1^H chemical shift in the bound and free state and ΔN is the subtraction of ^15^N chemical shift in the bound and free state. The overlapped residues in HSQC spectra were removed for the chemical shift perturbation calculation. Standard deviation multiplied by 4 was chosen as cut-off values (Williamson, 2013). Residues in interaction were removed from standard deviation calculation.

#### Ensemble Calculations

Ensembles of explicit models were generated using Flexible-Meccano (FM) (Ozenne et al., 2012b), which sequentially builds peptide planes based on amino acid specific conformational propensity and a simple volume exclusion term. To account for deviations from a random-coil description, different structure ensembles of 50,000 conformers were computed including user-defined local conformational propensities in different regions of the protein. Local conformational propensities were first localized using the consensus of all NMR data, and then were adjusted by comparing back-calculated and experimental ^1^DHN RDCs to get the lowest X^2^ (for more details (Senicourt et al., 2021)). For pm-β2AR-Cter, as for pm-β2AR-Cter (Senicourt et al., 2021), a long-range contact of 15 Å between two regions affected by the probe, i.e., from residues 338 to 357 and from residues 367 to 386 (regions in grey in Fig. S5), was introduced to get a better agreement between back-calculated and experimental ^1^DHN RDCs.

### FRET measurements

Fluorescence spectra were recorded on a Fluoromax-4 TCSP spectrofluorimeter (Horiba). Protein concentrations in the 0.5 μM range were used. Fluorescence emission spectra were recorded at 20 °C between 400 nm and 600 nm (λexc 343 nm) or 510 and 600 nm (λexc 500 nm) with 5 nm excitation and emission slits and 0.5 nm intervals. The absence of The FRET ratio corresponds to the ratio of the acceptor emitted fluorescence at 520 nm following excitation at two different wavelengths, 345 and 500 nm. FRET changes were calculated using the values obtained for arrestin in the presence of an irrelevant peptide as a reference and were expressed as %. Removal of acceptor bleedthrough and correction of direct acceptor excitation were carried out as described previously (Granier et al., 2007).

### Dissociation constant estimation

#### Microscale thermophoresis (MST)

Arrestin-2 was buffer-exchanged by using PD MiniTrap™ G-10 (Cytiva), in 50 mM Bis-Tris pH 7.5, 50 mM NaCl and 1 mM TCEP labeling buffer. Alexa Fluor® 488 Maleimide C5 (ThermoFisher Scientific) dye was added in 10-fold molar excess in 250 μL at 7 μM of arrestin-2. After 1 h at 4 °C, the reaction was quenched by adding 100 mM of L-cysteine and incubates during 15 min at room temperature. Buffer was exchanged and the free dye removed by gravity gel filtration using PD MiniTrap™ G-10 (Cytiva) into 50 mM HEPES pH 7.5, 50 mM NaCl. The labeling efficiency was calculated using absorbance at 280 nm (ε280 = 19,370 M^-1^.cm^-1^) for arrestin-2 and 495 nm (ε495 = 71,000 M^-1^.cm^-1^) for the dye. A correction factor of 0.11 was assumed for the absorbance at 495 nm to compensate the dye’s absorbance at 280 nm. After incubation of 5 nM of arrestin-2 labeled with an increasing amount of pm-GHSR-Cter (0.01-450 μM) at 20 °C during 30 min, sample were loaded into standard capillaries (Monolith NT Capillaries, Nanotemper Technologies). Before each measurement, all samples were centrifuged at 12,000 g during 15 min. Measurement was performed on a Monolith NT.115 MST apparatus (NanoTemper Technologies), by using a range excitation wavelength between 460 and 480 nm and a range emission wavelength measured between 515 and 530 nm. Dose-response curve was plotted using the MO.Affinity Analysis software (Nanotemper Technologies) and outlier values were removed. One-site binding model was used to calculate the Kd value.

#### Fluorescence anisotropy

pm-GHSR-Cter was buffer-exchanged by using PD MiniTrap™ G-10 (Cytiva), in labeling buffer (50 mM Bis-Tris pH 7, 50 mM NaCl and 1 mM TCEP). Alexa Fluor® 488 succinimidyl ester (ThermoFisher) was added in 10-fold molar excess in 100 μL at 50 μM of pm-GHSR-Cter. After an incubation of 1 h at room temperature, protected from light and under gentle agitation, the reaction was stopped by adding 10 μL of 1 M Tris and incubated for 15 min. Buffer was exchanged and the free dye removed by gravity gel filtration using PD MiniTrap™ G-10 (Cytiva) into labeling buffer. The labeling efficiency was calculated using absorbance at 280 nm (ε280 = 5500 M^-1^.cm^-1^) for pm-GHSR-Cter and 495 nm (ε495 = 71 000 M^-1^.cm^-1^) for the dye. A correction factor of 0.11 was assumed for the absorbance at 495 nm to compensate the dye’s absorbance at 280 nm. After 30 min incubation of 1 nM of pm-GHSR-Cter with increasing amounts of arrestin-2 (0-670 μM), samples were centrifuged at 12,000 g during 15 min. Measurement of fluorescence anisotropy was performed using a Tecan Safire II micro plate reader fluorescence spectrometer and a Corning 384 Low Flange Black Flat Bottom plate. Excitation and emission wavelengths were respectively measured at 470 nm and 530 nm. KD value was calculated by nonlinear regression analysis using a sigmoid curve fit from GraphPad Software

### Modeling of GPCR-Cter:Arrestin complexes

The complexes of GPCR-Cters and arrestin-2 were modeled using the following steps. First, the stretch of residues identified to be involved in binding based on NMR data was modeled in complex with arrestin-2 using the crystal structure of V2Rpp in complex with arrestin-2 (PDB ID: 4JQI). This included residues 356-364 for V2 and 360-367 for GHSR. These models were then refined using FlexPepDock server (**PMID: 21622962, 20455260**). The rest of the residues of Cters were added using the program RanCh (Bernado et al., 2007; Tria et al., 2015) followed by addition of backbone and side-chain atoms using PULCHRA (**PMID: 18196502**). These structures were then subjected to Molecular Dynamics (MD) simulations using the DES-amber force field (doi: 10.1021/acs.jctc.9b00251) obtained from https://github.com/paulrobustelli/Force-Fields. The simulation system was prepared using the tools available with gromacs 2020.5. This included addition of hydrogens, solvating in a box of TIP4P-D water molecules, neutralizing the system with the associated Na^+^ and Cl^-^ ions with scaled charges and setting the salt concentration to 0.15 M. We used periodic boundary conditions and restrained the bonds involving hydrogen atoms using LINCS algorithm (Hess et al., 1997). The electrostatic interactions were calculated using particle mesh Ewald (PME) (Darden et al., 1993) with a cutoff of 9 Å for long-range interactions. The system was minimized for 10,000 steps using steepest descent algorithm followed by NVT (constant number, volume and temperature) equilibration in 3 steps. In the first step, all the backbone atoms were restrained using a force constant of 1000 kJ/mol.nm^2^. In the second step, the region of Cters refined using FlexPepDock and the residues of arrestin-2 within 5 Å of this region were kept restrained while the rest of the proteins were allowed to move freely for 20 ns. In the third step, only the short region of the Cters, which forms beta-sheet, was restrained and the system was equilibrated for 20 ns. Finally, all the restraints were released and the 4 independent NPT (constant number, pressure and temperature) simulations of 250 ns each were carried out accumulating 1 μs of unrestrained simulations for each Cter. The temperature was maintained at 303.15 K using a Nose-Hoover thermostat (Nosé, 1984) and for NPT simulations, the pressure was maintained at 1 bar using Parrinello-Rahman barostat (Parrinello and Rahman, 1981).

### Accession numbers

The ^1^H, ^15^N and ^13^C assignment of pm-V2R-Cter, pm-GHSR-Cter and pm-β2AR-Cter variants are available in the BioMagResBank (http://www.bmrb.wisc.edu) under BMRB accession codes: **<u>51330</u>, <u>51328</u>, <u>51319</u>** respectively.

## Acknowledgements

We thank the synchrotron facility SOLEIL (St. Aubin) for allocating regular beam time (Block Allocation Group, proposal No. 20201085) and its dedicated staff for technical help with the beamline SWING. The authors are grateful to David Lemaire for MS support, and Alessandro Barducci and Matteo Paloni for simulated FRET efficiency differences.

## Funding

This work has benefited from support by the Labex EpiGenMed, an « Investissements d’avenir » program, reference ANR-10-LABX-12-01 awarded to MG.. N.S. and J.L.B. acknowledge the support of the ANR GPCteR (ANR-17-CE11-0022-01). This work was granted access to the HPC resources of CALMIP supercomputing center under the allocation 2022A-P22037 to N.S. The CBS is a member of the French Infrastructure for Integrated Structural Biology (FRISBI) supported by the French National Research Agency (ANR-10-INSB-05).

## Credit author statement

Investigation, methodology, writing-original draft preparation, M.G. and A.M.; Investigation and Resources, A.F. and F.A.; Formal analysis, A.S., A.T. and P.B.; Investigation, writing-review and editing, J.-L.B.; Conceptualization, supervision, funding acquisition, writing-original draft, review and editing, N.S. All authors have read and agreed to the published version of the manuscript.

## Additional files

Supplementary files

## Conflict of interest

The authors declare no conflict of interest.

## Abbreviations

IDP: intrinsically disordered protein
IDR: intrinsically disordered region
PTM: post-translational modifications
GPCR: G protein-coupled receptor
GRK: G protein receptor kinase
V2R: vasopressin V2 receptor
GHSR: ghrelin receptor 1a
β2AR: β2-adreneric receptor
wt-V2R-Cter: wild-type V2R C-terminus
pm-V2R-Cter: phosphomimetic V2R C-terminus
wt-GHSR-Cter: wild-type GHSR C-terminus
pm-GHSR-Cter: phosphomimetic GHSR C-terminus
wt-β2AR-Cter: wild-type β2AR C-terminus
pm-β2AR-Cter: phosphomimetic β2AR C-terminus
V2Rpp: synthetic fully phosphorylated peptide of V2R C-terminus
C7pp: synthetic fully phosphorylated peptide of CXCR7 C-terminus
pRho-Cter: phosphorylated C-terminus of rhodopsin receptor
PPII: polyproline helix 2
CD: circular dichroism
MS: mass spectrometry
MST: microscale thermophoresis
FRET: fluorescence resonance energy transfert
SAXS: small angle X-ray scattering
SDS-PAGE: sodium dodecyl sulphate-polyacrylamide gel electrophoresis
SEC-MALS: size exclusion chromatography – multi angle light scattering
NMR: nuclear magnetic resonance
SCS: secondary chemical shift
CSP: chemical shift perturbation
HSQC: heteronuclear single quantum coherence
NOE: nuclear Overhauser effect
PRE: paramagnetic relaxation enhancement
RDC: residual dipolar coupling
FM: flexible meccano
EDTA: EthyleneDiamineTetraAcetic acid
DTT: dithiothreitol
Cou: L-(7-hydroxycoumarin-4-yl ethylglycine
PMSF: phenylmethylsulfonyl fluoride
TCEP: Tris(2-CarboxyEthyl)Phosphine

---

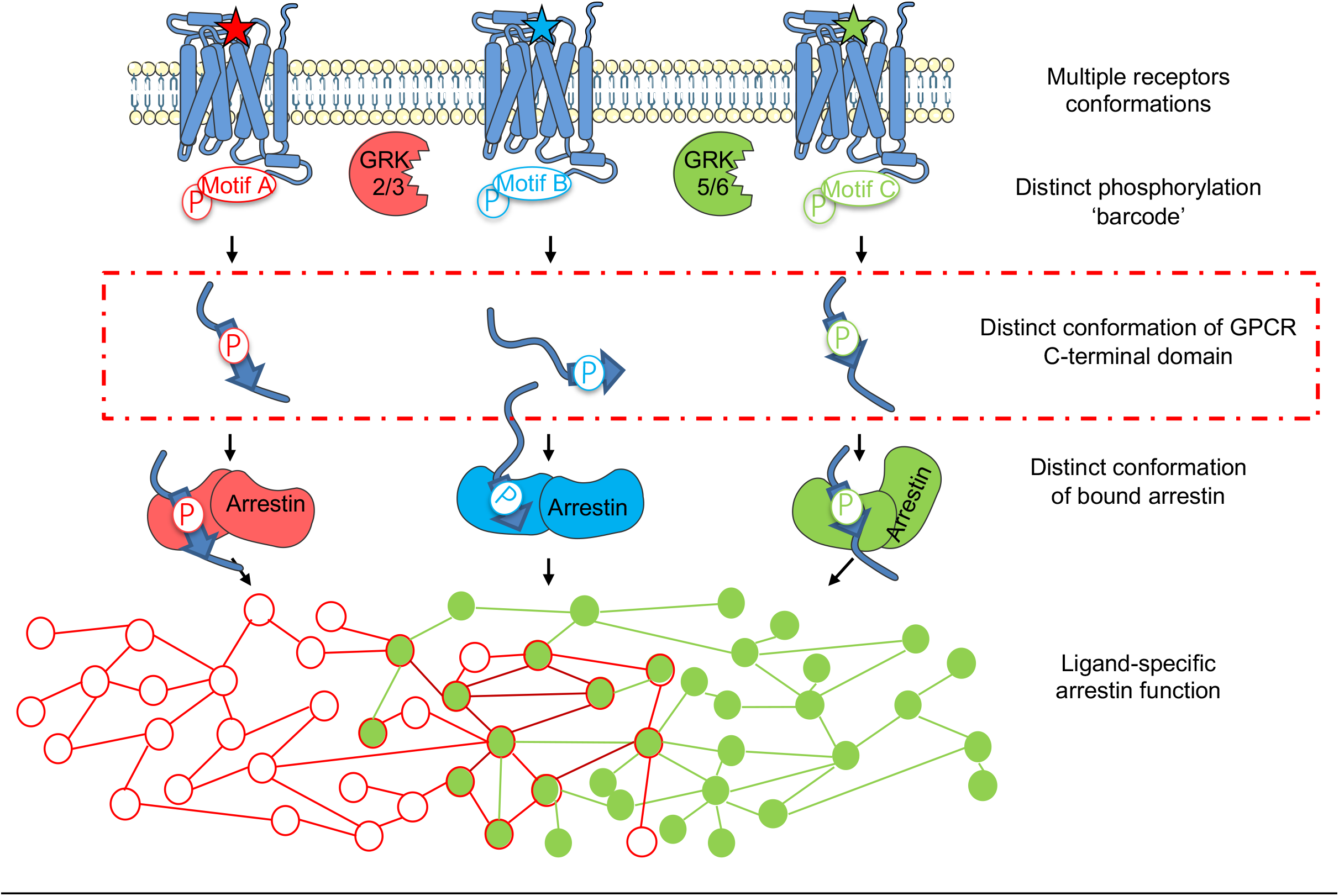

## Notes

### Competing Interest Statement

The authors have declared no competing interest.

