## Supplementary Information for "Phosphorylation motif dictates GPCR C-terminal domain conformation and arrestin interaction"

| Protein | 3D | $^3J_{HNHA}$ | Relaxation | RDC alcohol | RDC Pf1 | PRE |
|---|---|---|---|---|---|---|
| pm-V2R | 50 | 93 | 93 | 93 | 138 | NT |
| pm-GHSR | 377 | 263 | 263 | 263 | 250 | NT |
| pm-β2AR | 206 | 476 | 476 | 476 | 212 | 100 |

**Table S1 Concentration used for the NMR characterization of phosphomimetic GPCR C-termini of V2R-Cter (blue), GHSR-Cter (green) and β2AR-Cter (pink).** Concentration used in each experiment is indicated in μM. The letters NT mean that these experiments have not been carried out.

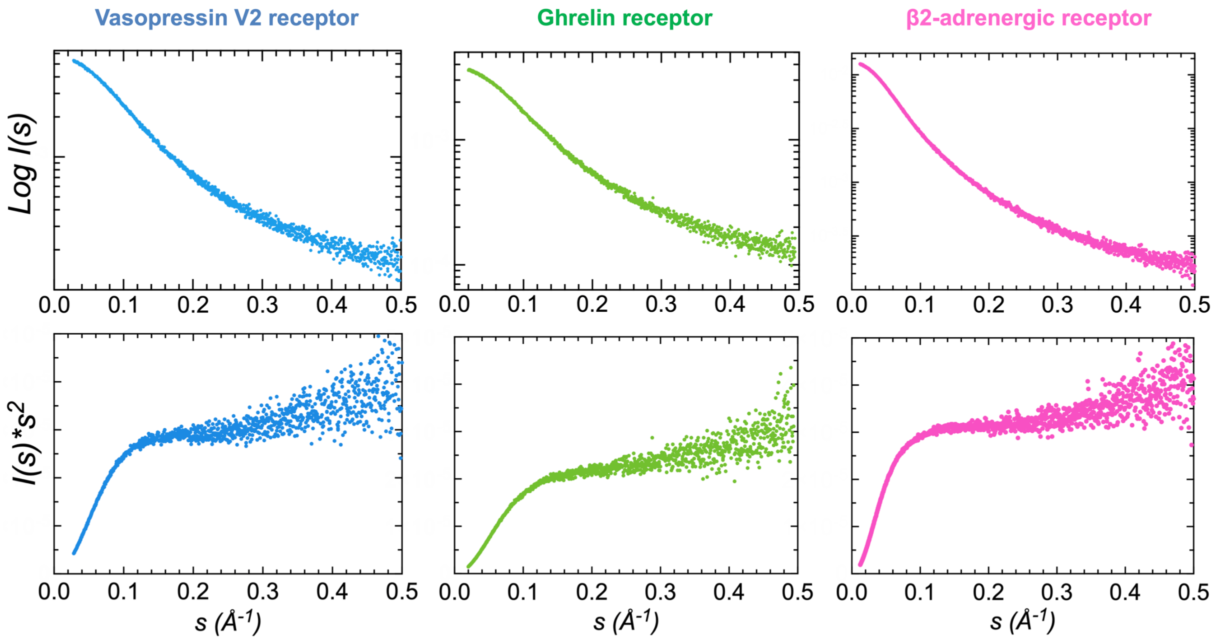


**Fig. S1 Related to Fig. 1 SAXS data of phosphomimetic variants of V2R-Cter (blue, left), GHSR-Cter (green, middle) and β2AR-Cter (pink, right).** Semi-logarithmic representation of the experimental SAXS curves versus scattering angle (upper panel) and Kratky plots (lower panel). The Kratky plot of variants display a typical profile of IDP with no clear maximum and a monotone increase along the momentum transfer range[1–3].

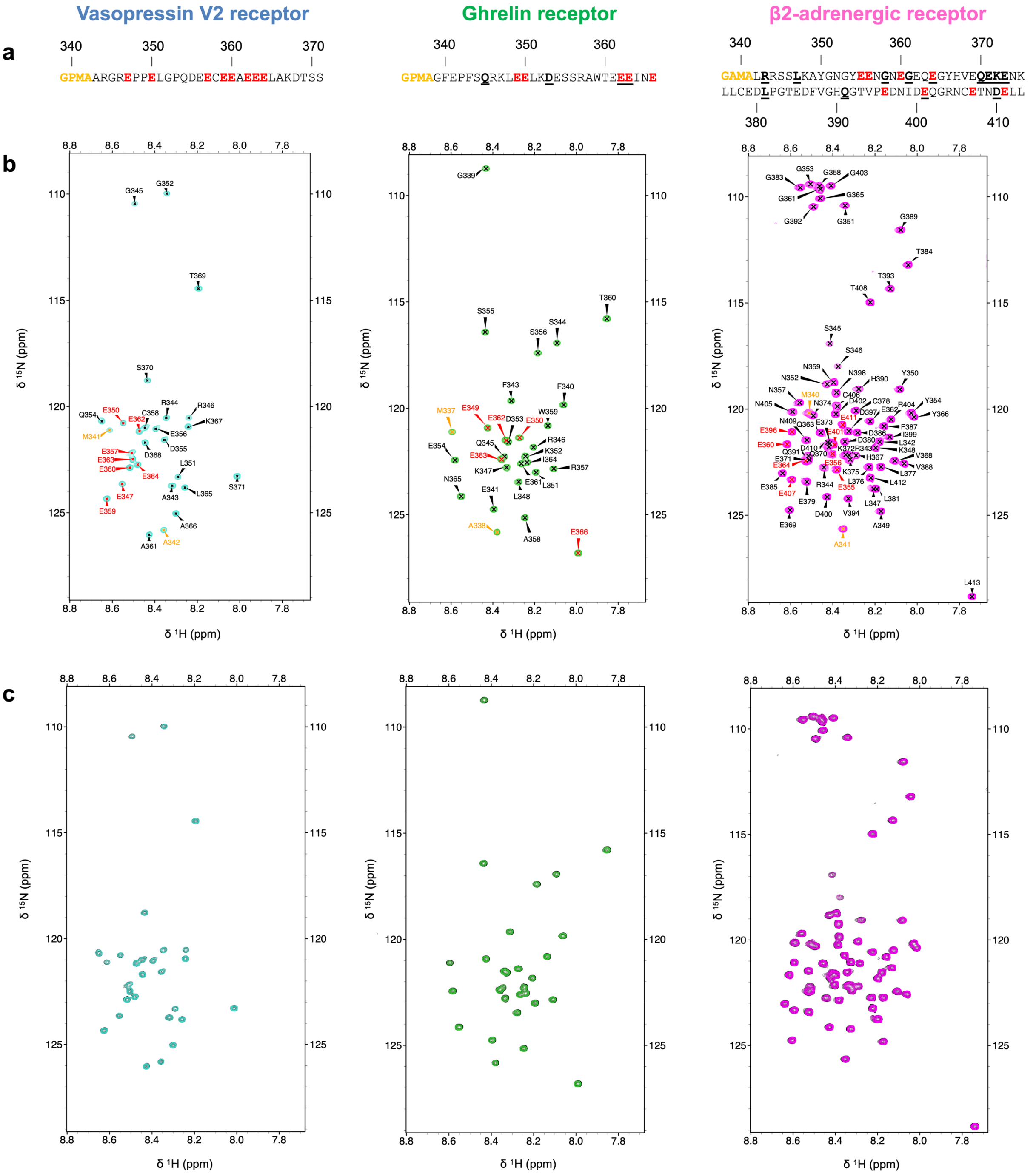


**Fig. S2 Related to Fig. 2b Assignments of phosphomimetic variants of V2R-Cter (blue, left), GHSR-Cter (green, middle) and β2AR-Cter (pink, right). a** Sequence of the phosphomimetic C-terminal domains with known phosphorylation sites mutated by glutamic acid (E) indicated in red. Amino acids coming from the protease cleavage site are shown in orange and overlapping residues are underlined in black. **b** $^{15}N$-HSQC of the C-terminal domains were recorded on 300 µM samples at 700 MHz, 20 °C, in 50 mM Bis-Tris pH 6.7, 150 mM NaCl buffer. Residue-specific assignments are shown on spectra and are colored according to **a**. **c** Overlay of $^{15}N$-HSQCs recorded on samples at 300 µM (black) or 100 µM (colored) at 700 MHz, 20 °C, 50 mM Bis-Tris pH 6.7, 150 mM NaCl of V2R-Cter (blue, left), GHSR-Cter (green, middle) and β2AR-Cter (pink, right). No chemical shift changes were observed in NMR spectra at different concentrations. Thus, the spectrum at 300 µM (black) is hidden by the spectrum at 100 µM (colored). The concentration of pm-GPCR-Cters doesn't affect their conformations.

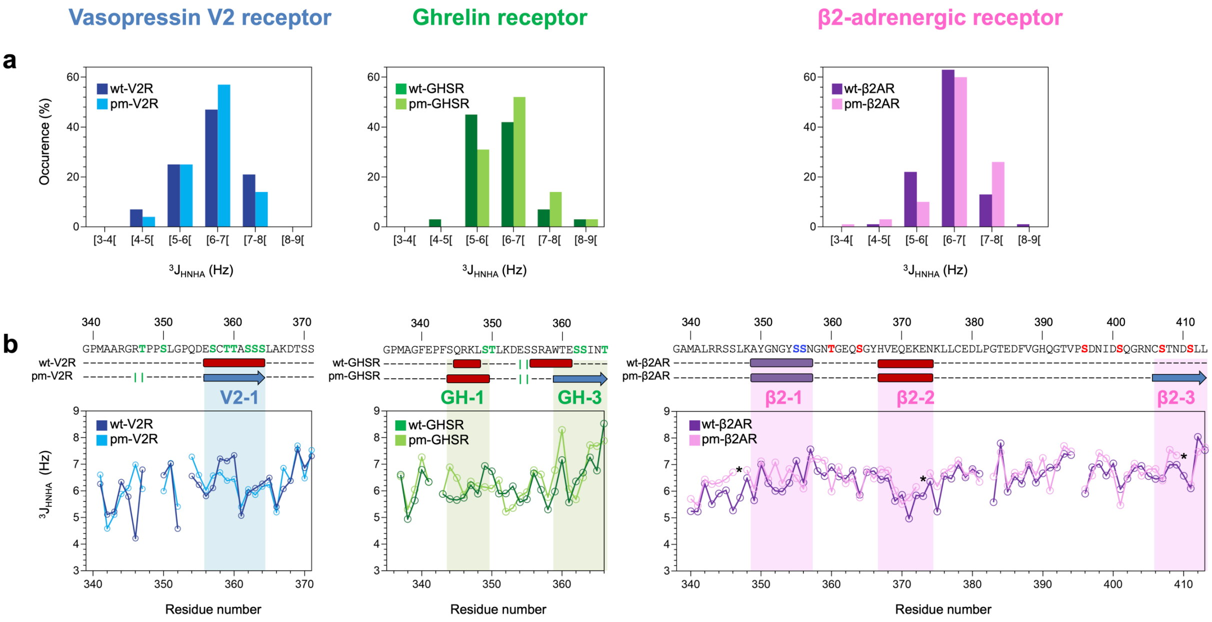


**Fig. S3 Related to Fig. 2 Overlay of 3JHNHA scalar coupling of wild type (dark color, data previously shown [4]) and phosphomimetic variant (light color) form of V2R-Cter (blue, left); GHSR-Cter (green, middle) and β2AR-Cter (purple, right). a** Histogram of the 3JHNHA in Hz. The couplings seem to shift to higher values in the variant suggesting a lower content of helical conformation (values below 6 Hz) and a higher content of extended conformation (values above 8 Hz). **b** On top, sequence and NMR secondary structure consensus of GPCR C-termini are shown according to figure 2. Below, the coupling values along the sequence are represented. For β2AR variant, outlier values for overlapping residues in HSQCs were removed (dark stars). They were recorded at 700 MHz, 20 °C, 50 mM Bis-Tris pH 6.7, 150 mM NaCl.

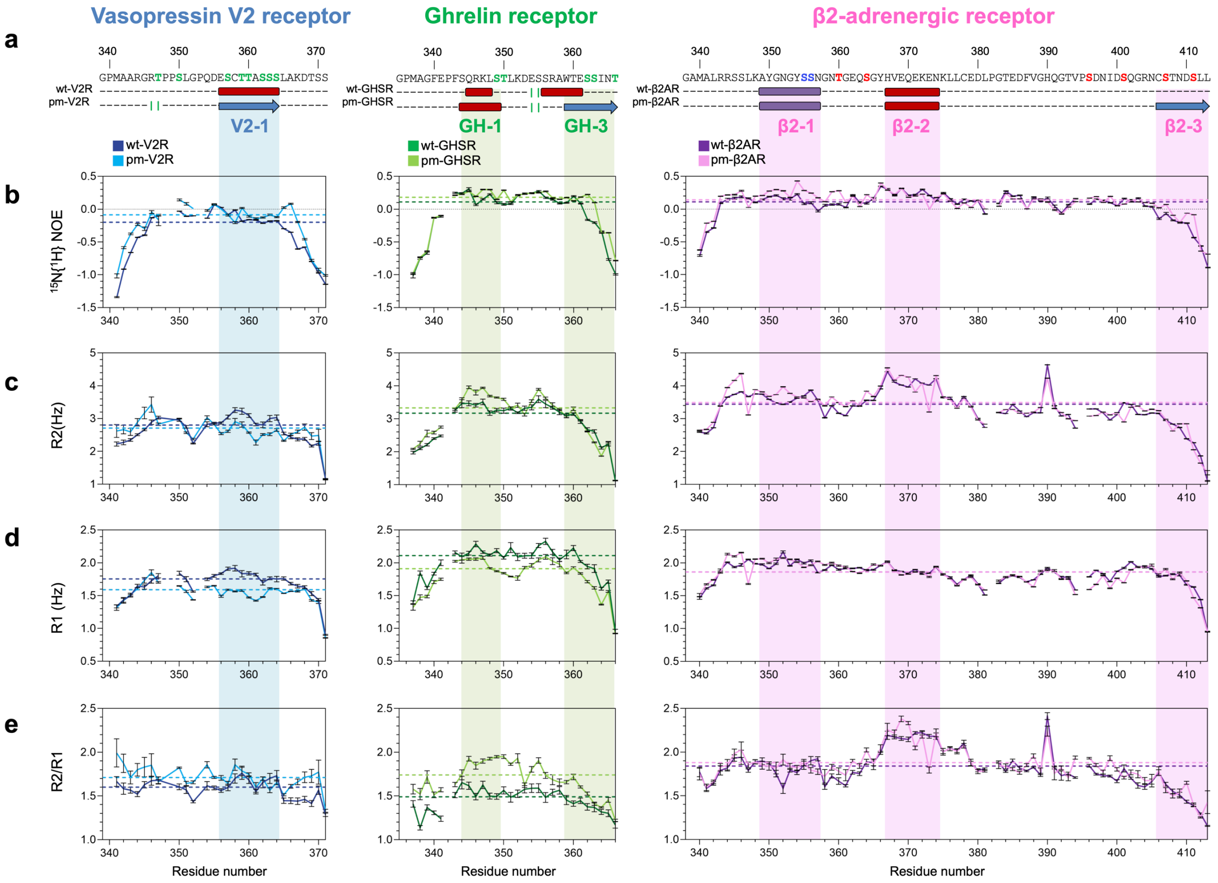


**Fig. S4 Related to Fig. 2c Overlay of relaxation data of wild type (dark color, data previously shown [4]) and variant (light color) forms of V2R-Cter (blue, left); GHSR-Cter (green, middle) and β2AR-Cter (purple/pink, right). a** Sequence and NMR secondary structure consensus of GPCR C-termini are shown according to Fig. 2. **b** $^{15}N\{^{1}H\}$ heteronuclear NOE. **c** Transversal relaxation rates R2. **d** Longitudinal relaxation rates R1; and **e** R2/R1 ratio are shown in colored line. Average values are indicted by dashed colored line. Relaxation data were recorded at 700 MHz, 20 °C, in 50 mM Bis-Tris pH 6.7, 150 mM NaCl buffer.

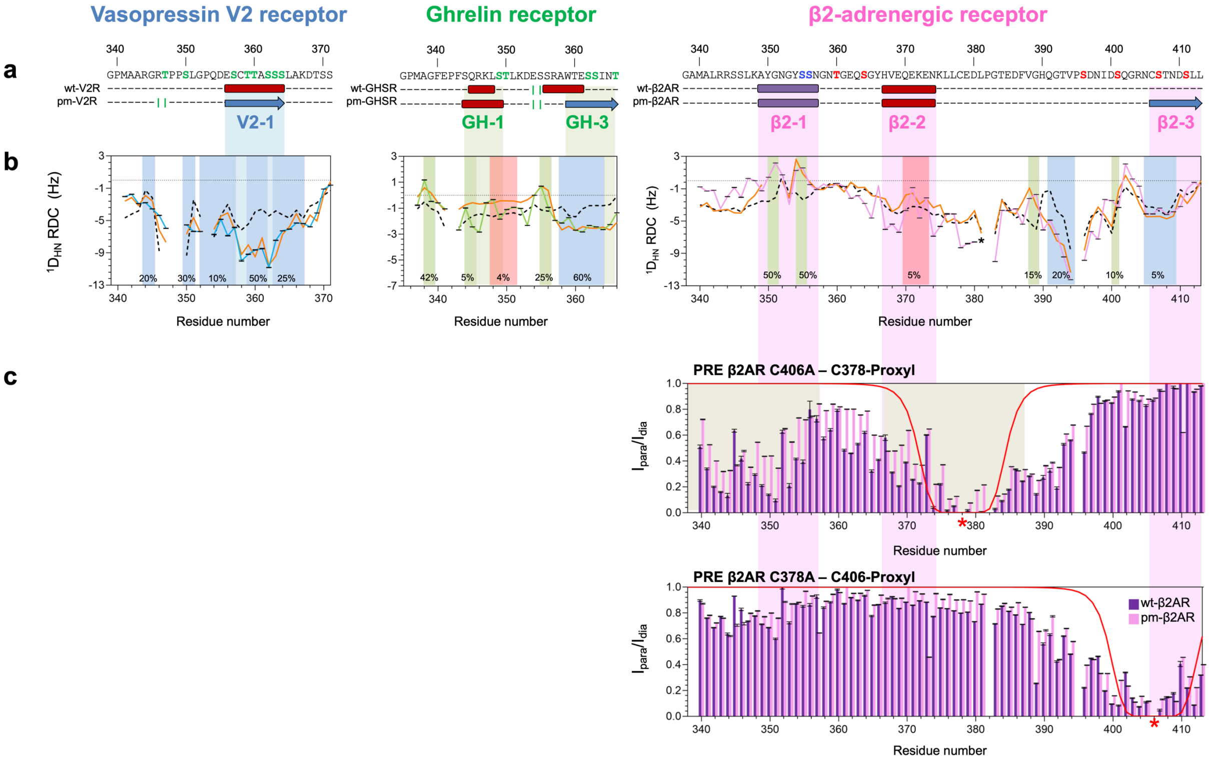


**Fig. S5 Related to Fig. 2d $^1D_{NH}$ RDCs values and PRE data used in the ensembles description of wild type (dark color, data previously shown [4]) and phosphomimetic variant (light color) forms for V2R-Cter (blue, left); GHSR-Cter (green, middle) and β2AR-Cter (purple/pink, right). a** Sequence and NMR secondary structure consensus of GPCR-Cters are shown according to Fig. 2. **b** Comparison of experimental RDCs values (full line) measured in bacteriophage Pf1 for pm-GHSR-Cter, and alcohol mixture for pm-V2R-Cter and pm-β2AR-Cter with back-calculated RDC computed by FM on random-coil (black dash line), or on biased ensemble (orange line) populated as indicated in the figure (regions colored in red (helical structure), blue (extended structure) or green (turn). Outliers due to overlapping residues were removed (dark star). **c** PRE data of C406A variant labeled on C378 (red star). Reduction of the intensity ratio suggests long-range interaction in regions highlighted in grey, which has been used for FM building of pm-β2AR-Cter and wt-β2AR-Cter[4] ensembles. PRE data of C378E variant labeled on C406 (red star). PREs were calculated from the ratio of intensity ($I_{para}/I_{dia}$) of $^{15}$N-HSQC spectra measured for the paramagnetic ($I_{para}$) sample (position of PROXYL probe is indicated with a red star), and for the diamagnetic sample ($I_{dia}$). They were recorded at 700 MHz, 20 °C, in 50 mM Bis-Tris pH 6.7, 150 mM NaCl buffer, on sample at 50 µM for wt-β2AR-Cter and 100 µM for pm-β2AR-Cter. Theoretical PRE curves for a completely disordered protein are shown in red.

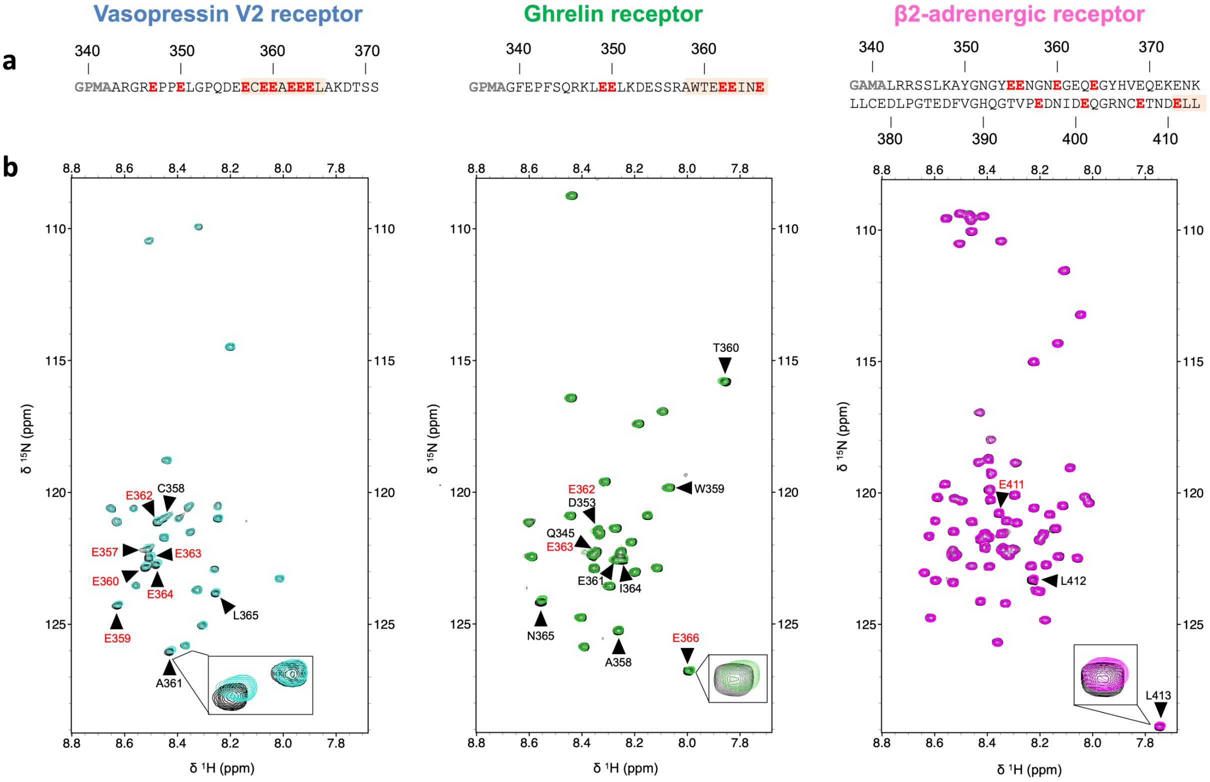


**Fig. S6 Related to Fig. 3 Visualization of pm-GPCR-C-ter:arrestin interaction by NMR of V2R-Cter (blue, left), GHSR-Cter (green, middle) and β2AR-Cter (pink, right). a** Sequence of the GPCR C-termini. Glutamic acid residues that mimic GRK phosphorylation sites are indicated in red. Binding regions are highlighted in orange. **b** Overlay of $^{15}$N-HSQC spectra without arrestin (in black) and with arrestin at ratio 1:10 (in color). Residues involved in the interaction are noted in black or red according to **a**. Spectra were recorded on $^{15}$N labeled Cter samples at 10 µM with 0 or 100 µM of unlabeled arrestin at 800 MHz, 20 °C, 50 mM Bis-Tris pH 6.7, 50 mM NaCl.

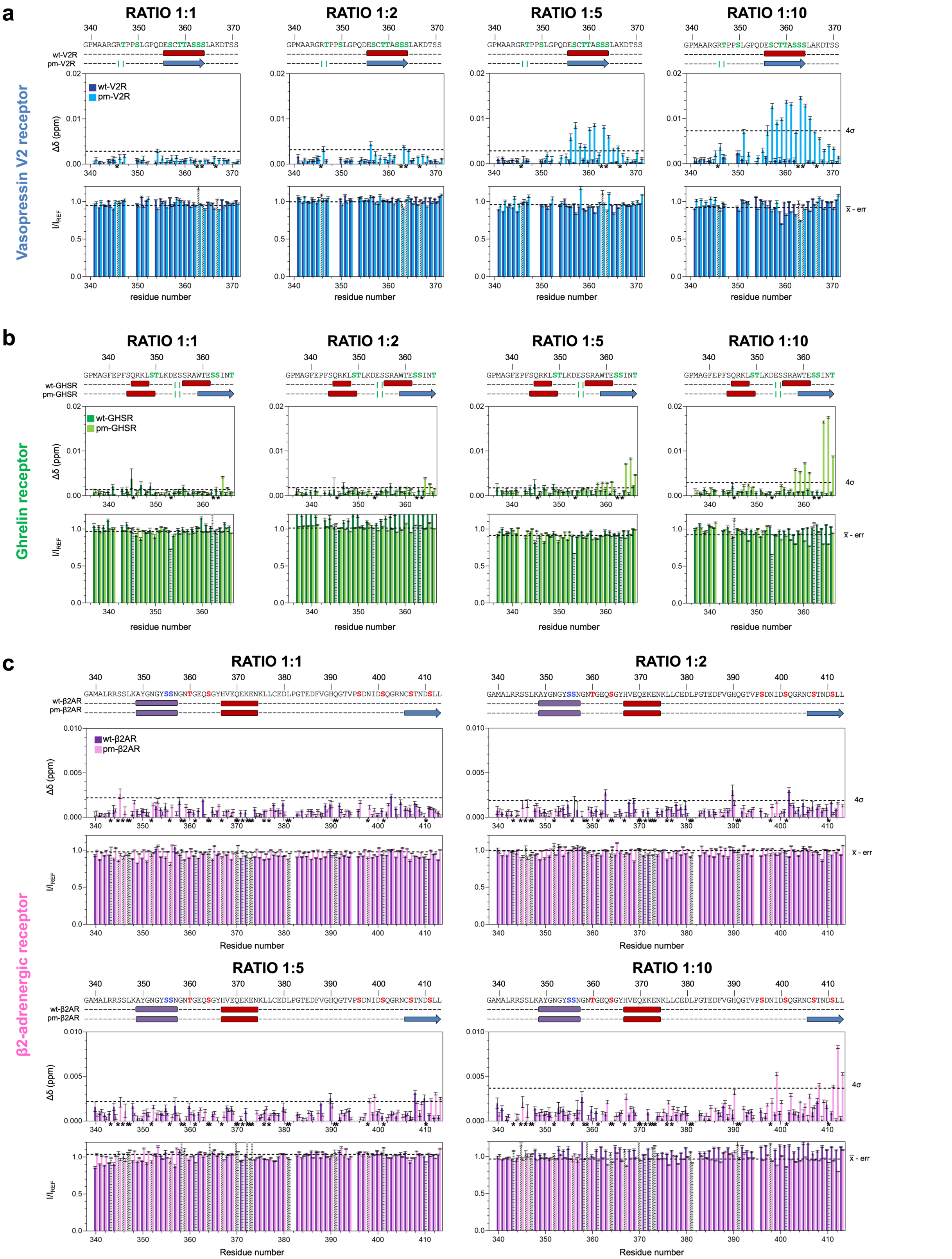


**Fig. S7 Related to Fig. 3 Interaction by NMR between arrestin-2 and (a) wt-V2R-Cter (dark blue) or pm-V2R-Cter (light blue), (b) wt-GHSR-Cter (dark green) or pm-GHSR-Cter (light green) and (c) wt-β2AR-Cter (purple) or pm-β2AR-Cter (pink) at different ratios.** For each C-terminus, the sequence and NMR secondary structure consensus are indicated according to Fig. 2. Below, histogram of $^{1}H/^{15}N$ chemical shift changes and intensity ratios ($I/I_{REF}$) are shown. Cut-off are indicated in black, dashed line and on the right. In the histogram of chemical shit changes, overlapping residues in HSQCs were removed and are shown in black stars near residues numbers. In the histogram of intensity ratio, they are represented in dashed bars. Spectra were recorded at 800 MHz, 20 °C, 50 mM Bis-Tris pH 6.7, 50 mM NaCl, on $^{15}N$ Cter samples at 10 µM C-termini with 0, 10, 20, 50 or 100 µM of unlabeled arrestin.

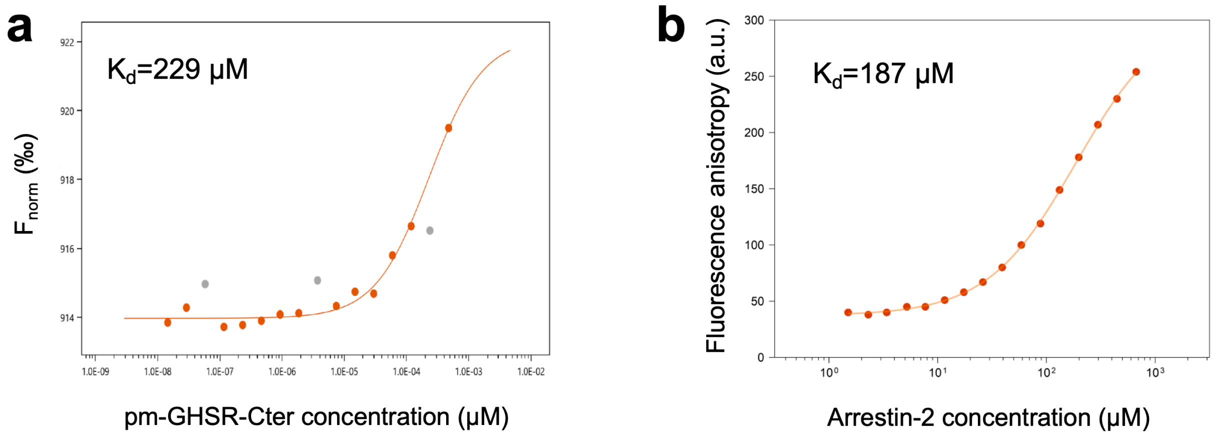


**Fig. S8 Low affinity interaction between arrestin-2 and the C-terminus of GHSR phosphomimetic. a** Dose-response curve of the Micro-Scale Thermophoresis (MST) titration. Labeled arrestin-2 at 5 nM was incubated with GHSR peptide (from 0.01 to 450 μM), at pH 6.7, 20 °C, 50 mM NaCl. An MST on-time of 5 sec was used for analysis and a dissociation constant ($k_D$) of 229 μM was obtained. Three data points were removed from the calculation (grey dots). **b** Fluorescence anisotropy curve. Labeled GHSR was incubated with varying amounts of arrestin-2: from 0 to 670 μM, at pH 6.7, 20 °C, 50 mM NaCl. A 187 μM dissociation constant ($k_D$) was determined.

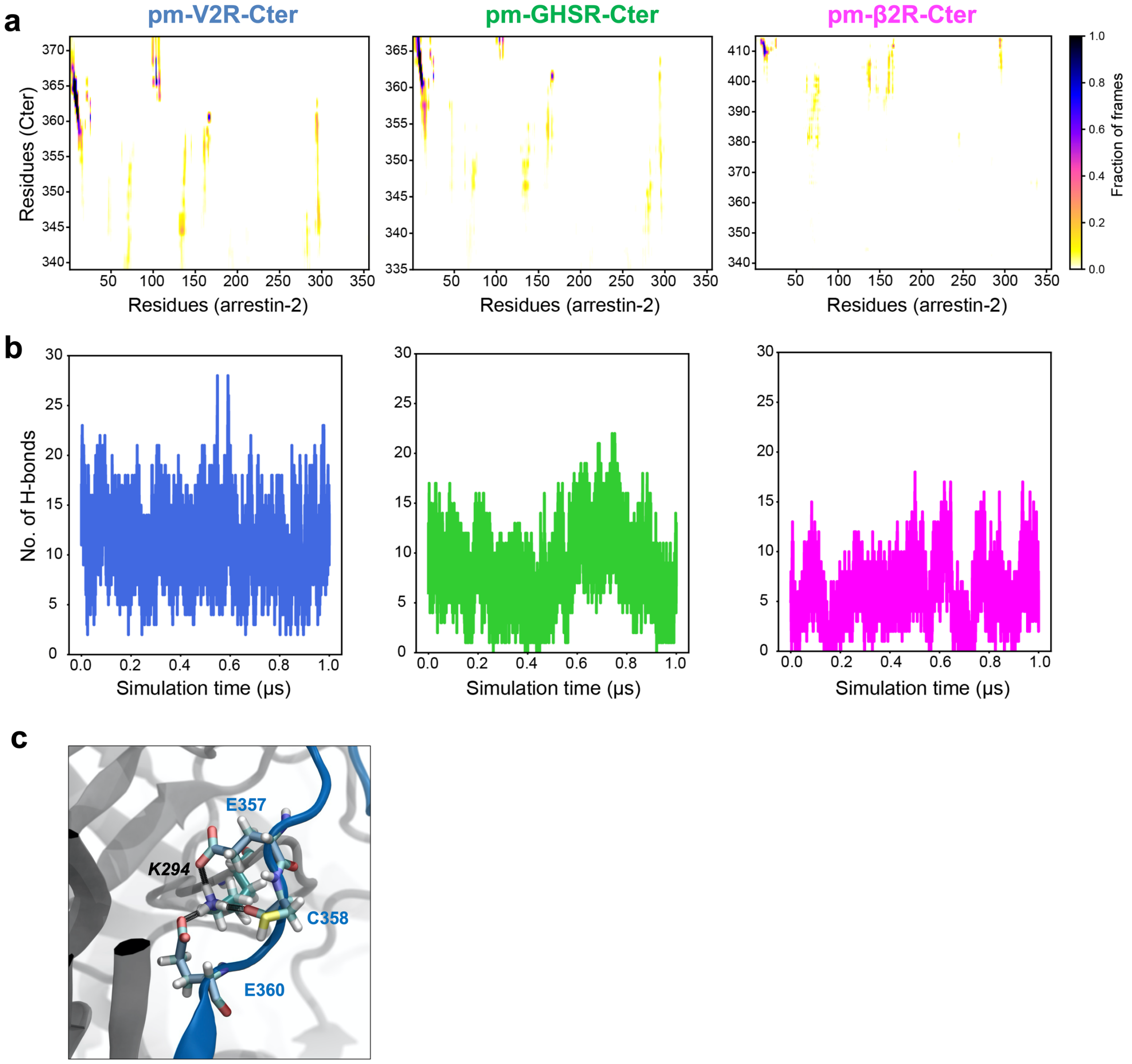


**Fig. S9 Interaction analysis of pm-V2R-Cter (blue), pm-GHSR-Cter (green) and pm-β2AR-Cters (pink): arrestin (grey) modeling complexes. a** The lifetime of contacts between residues of arrestin-2 (X-axis) and C-ters (Y-axis). **b** The number of hydrogen bonds between the Cters and arrestin through the concatenated simulations trajectories. **C** The bifurcated salt-bridge formed by E357 and E360 of pm-V2R-Cter with K294 of arrestin-2 along with an additional H-bond with C358 (pm-V2R-Cter)

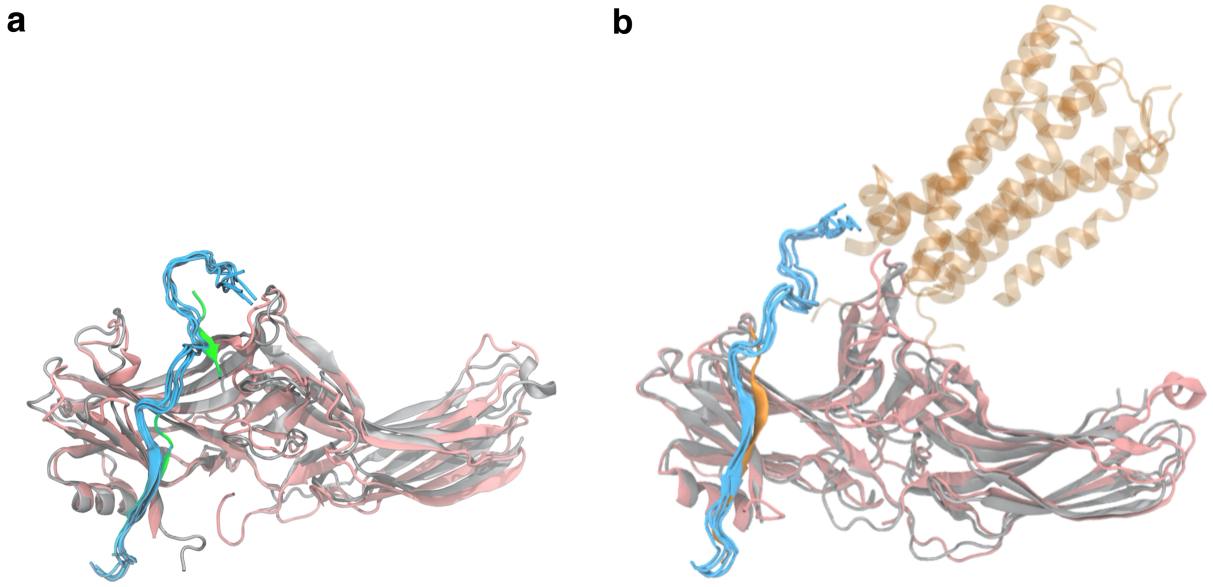


**Fig S10. Atomic structures of pm-V2R-Cter and entire V2 receptor complexed with arrestin.** Four selected peptide conformation from an ensemble obtained from MD simulations of pm-V2R-Cter (light blue) complexed to arrestin (pink) superimposed to : **a** the X-ray structure of the V2Rpp (green, PDB code 4JQI) and to **b** the cryo-EM structure of the entire V2 receptor (orange, PDB code 7R0J), complexed to arrestin (grey).
